# Optogenetics in *Sinorhizobium meliloti* enables spatial control of exopolysaccharide production and biofilm structure

**DOI:** 10.1101/2020.07.03.187138

**Authors:** Azady Pirhanov, Charles M. Bridges, Reed A. Goodwin, Yi-Syuan Guo, Jessica Furrer, Leslie M. Shor, Daniel J. Gage, Yong Ku Cho

**Affiliations:** Department of Biomedical Engineering, University of Connecticut, Storrs CT, USA; Department of Molecular and Cellular Biology, University of Connecticut, Storrs CT, USA; Department of Chemical and Biomolecular Engineering, University of Connecticut, Storrs CT, USA; Department of Computer Science, Physics, and Engineering, Benedict College, Columbia, SC, USA; Center for Environmental Sciences and Engineering, University of Connecticut, Storrs CT, USA; Institute for Systems Genomics, University of Connecticut, Storrs CT, USA

**Keywords:** soil bacteria, exopolysaccharide, synthetic biology, optogenetics, biofilm

## Abstract

Microorganisms play a vital role in shaping the soil environment and enhancing plant growth by interacting with plant root systems. Due to the vast diversity of cell types involved, combined with dynamic and spatial heterogeneity, identifying the causal contribution of a defined factor, such as a microbial exopolysaccharide (EPS), remains elusive. Synthetic approaches that enable orthogonal control of microbial pathways are a promising means to dissect such complexity. Here we report the implementation of a synthetic, light-activated, transcriptional control platform in the nitrogen fixing soil bacterium *Sinorhizobium meliloti.* By fine tuning the system, we successfully achieved optical control of an EPS production pathway without significant basal expression under non-inducing (dark) conditions. Optical control of EPS recapitulated important behaviors such as a mucoid plate phenotype and formation of structured biofilms, enabling spatial control of biofilm structures in *S. meliloti.* The successful implementation of optically controlled gene expression in *S. meliloti* enables systematic investigation of how genotype and microenvironmental factors together shape phenotype *in situ*.

**Significance:** Microorganisms are key players in sustaining the soil environment and plant growth. Symbiotic associations of soil microbes and plants provide a major source of nitrogen in agricultural systems, prevent water contamination from synthetic fertilizer application, and support crop growth in marginal soils. However, measuring the impact of microbial gene products on beneficial function remains a major challenge. This work provides a critical step toward addressing this challenge by implementing external gene regulation in a well characterized nitrogen fixing soil bacterium. We show that light exposure enables spatial and temporal control of the extracellular polysaccharide production functionality essential for symbiosis. Remote control of genes enables the benefits of candidate microorganisms to be systematically measured and enhanced within complex natural settings.

## Introduction

The soil bacterium *S. meliloti* is a Gram-negative α-proteobacterium capable of fixing atmospheric nitrogen during a symbiotic association with certain host plants such as alfalfa (*Medicago sativa*). Rhizobial nitrogen fixation through symbiotic associations with crop legumes such as soybean, oilseed legumes, chickpea, and common beans is the most important source of natural fixed nitrogen in agricultural systems (1, 2). This symbiosis contributes significantly to sustainable agriculture by reducing water contamination from nitrogenous fertilizers (3) and supports crop growth in marginal soils of arid regions (3, 4). *S. meliloti* is one of the most thoroughly studied rhizobia, with complete genome sequences analyzed for multiple isolates (5–8), including their megaplasmids that carry genes of critical physiological and symbiotic importance (9). *S. meliloti* is related to plant pathogens such as *Agrobacterium* and animal pathogens such as *Brucella. S. meliloti* and its pathogenic relatives are important models for studying host-microbe interactions (8).

Exopolysaccharides (EPS) are produced by a wide range of bacteria, and impart many physiological functions including biofilm structure, nutrient acquisition, environmental stress resistance, and resistance to antimicrobials (10–12). EPS produced by rhizobia have been extensively studied due to their essential role in host plant invasion (13–15). Lab strains of *S. meliloti* produce succinoglycan (EPS I) and galactoglucan (EPS II) through coordinated activity of genes located on the pSymB megaplasmid and the chromosome (9, 16–20). EPS I biosynthesis involves the *exo/exs* genes, which are organized together in several operons on pSymB. Mutations within a number of these genes result in complete abolition of EPS I production (18, 19, 21,22). EPS II biosynthesis requires the *exp* gene cluster, which is organized into five putative transcriptional units, among which *wga (expA), wgcA (expC), wgd (expD*) and *wge (expE*) contain structural genes that are needed for the production of EPS II (20, 23–25).

The control of EPS synthesis in *S. meliloti* involves multiple regulatory systems (23). Environmental cues, nutrient availability, osmolarity, and the ionic strength of the surrounding medium affect the production of EPS in *S. meliloti* (26–34). In particular, low phosphate levels, commonly encountered in soil, are sensed by the two-component regulatory system PhoR-PhoB, which stimulates EPS II production (24, 35, 36). EPS II production is also coupled to quorum sensing and motility via regulatory proteins MucR, ExoR and ExpR (23, 37, 38).

EPS production is influenced by environmental factors, yet EPS production also indirectly influences these same environmental factors by altering the structure and function of rhizosphere systems. Regarding its nitrogen fixing functionality, EPS biosynthesis is required for invasion of plant roots to occur (15, 16, 39, 40). In addition, EPS promotes auto-aggregation of planktonic *S. meliloti* cells and the formation of structured biofilms (30, 41). Moreover, recent results suggest that EPS affects how cells interact with the soil particle surfaces and affects water retention and resiliency in soil, as shown with emulated soil micromodels (42–44). The ability to spatially and temporally control EPS biosynthesis apart from its multitude of endogenous regulatory pathways is a powerful new tool to assess not only the causal role of EPS in these and other processes but in general to decode how genotype gives rise to phenotype *in situ.*

Here we report an implementation of an orthogonal, synthetic gene expression system in *S. meliloti* for controlling EPS II biosynthesis. We show that the blue-light activated transcription factor EL222 from *Erythrobacter litoralis* HTCC2594 is functional in *S. meliloti* and can be used to regulate gene expression. Characterization of synthetic ribosome binding sites and alternative stop codons enabled engineering of an optically controlled expression platform with minimal expression in the absence of blue light. By optically controlling expression of EPS II biosynthesis genes we demonstrated spatial control of structured biofilm formation. The successful implementation of an optically controlled gene expression system in *S. meliloti* opens the door to testing the function of EPS genes, and as well as many others, *in situ.*

## Methods

### Growth conditions

*E. coli* strain TOP10 (Invitrogen) was grown in LB medium or on LB plates at 37 °C for plasmid preparation. Antibiotics were used for *E. coli* cultures at the following concentrations (μg mL^-1^): tetracycline, 10; gentamycin, 15. *S. meliloti* was grown in, TY medium (6 g of tryptone, 3 g of yeast extract, and 0.38 g of CaCl2 per liter), M9 medium (5.8 g Na_2_HPO_4_, 3 g KH_2_PO_4_, 0.5 g NaCl, 1 g NH_4_Cl, 0.5 mg biotin, 0.011 g CaCl_2_, 0.12 g MgSO_4_, 4 g glucose per liter), or Rhizobium Defined Medium (RDM) (0.6 g KNO_3_, 0.1 g CaCl_2_·2H_2_O, 0.25 g MgSO_4_7H_2_O, 0.01 g FeCl_3_·6H_2_O, 0.5 mg biotin, 0.01 g thiamine, 20 g sucrose, 1 gram each of K_2_HPO_4_ and KH_2_PO_4_ per liter). For low-phosphate RDM, 1:1 mass ratio of K_2_HPO_4_:KH_2_PO_4_ was used at a final total concentration of 0.1 mM and morpholinepropanesulfonic acid (MOPS) was added as a buffer at a final concentration of 0.01 g mL^-1^ and pH of 7.0. Unless otherwise stated, *S. meliloti* cultures were grown in 5 mL TY at 30 °C shaken at 230 RPM and media were supplemented with antibiotics at the following concentrations (μg mL^-1^): streptomycin, 500; gentamycin, 30; tetracycline, 5. Dark cultures were grown in culture tubes wrapped with aluminum foil.

Standard electroporation technique was used to transform *S. meliloti* cells (45). Briefly, 50 μL of electro-competent *S. meliloti* cells were mixed with 1.5 μL of DNA (~50 ng total) in sterile Eppendorf tubes chilled on ice. Subsequently, cells were transferred to ice chilled 0.1 cm electrode gap Gene Pulser Cuvettes (Bio-Rad cat #1652089) and an electric pulse of 2.3 kV was applied using GenePulser XCell (Bio-Rad). 1 mL of SOC medium was added and transformants were selected on TY plates with the appropriate antibiotics.

For making frozen culture stocks, cells were grown for 2 d at 30 °C. Following the incubation, aliquots of 20% glycerol stock cultures were prepared by mixing equal volume of culture with filter sterilized 40% glycerol (v/v in water) solution and stored at −80°C. When necessary one vial of stock cell culture was then thawed on ice and 20 μL was inoculated in 3 mL of TY medium with appropriate antibiotics.

### Construction of *S. meliloti* EPS I and EPS II deletion strains

*S. meliloti* strain Rm8530 was used as the parent to construct isogenic, unmarked, mutants with deletions in genes required for synthesis of EPS I (RG27; Δ*exoY*), EPS II (RG33; Δ*wgaAB*) and a double mutant unable to make either (RG34; Δ*exoY*Δ*wgaAB*). In addition, the RG33 (Δ*wgaAB*) and the double mutant RG34 (Δ*exoY*Δ*wgaAB*) strains were complemented with an integrated plasmid, pRG73 (*wgaAB^+^*) that expressed the *wgaAB* genes from their native promoter. A detailed procedure of *S. meliloti* strain construction is described in **Supp. Methods**.

### Plasmid construction

Plasmids and primers used in this study are described in **Table 1** and **Supp. Table 2**. Superfolder GFP (*sfgfp*) (46) was cloned under P_El222__*rbsD* promoter-ribosome binding site (RBS) combination in pSEVA531 by using Golden Gate (GG) assembly. The resulting plasmid was named pAP01. Next, the *El222* gene driven by BBa_J23105-rbs34 and BBa_J23115-rbs34 promoter-RBS combinations were cloned into the pAP01 backbone and the resulting plasmids were named pAP05 and pAP15 respectively. The *wgaAB* genes were PCR amplified from wild type *S. meliloti* genomic DNA and a P_El222__*rbs34_wgaAB* insert was assembled. Subsequently, P_El222__*rbsD_sfGFP* in pAP05 was swapped with P_El222__rbs34_*wgaAB* using GG assembly, which produced plasmid pAP14, where *El222* expression was driven by the BBa_J23105-rbs34 promoter-RBS combination. In order to construct plasmids pAP33, pAP34, pAP35, pAP36, pAP37 and pAP38 which have different combinations of ribosome binding site and start codon, components were engineered with flanking XbaI and NotI sites. Briefly, the pAP05L backbone (which constitutively expressed *EL222* from *lacI^q^* promoter) was PCR amplified with a reverse primer containing a 5’ XbaI site flanking a ribosome binding site (either rbsD, rbs31 or rbs33) and forward primer containing a NotI restriction site. The *wgaAB* gene was PCR amplified with primers containing either ATG or GTG as a start codon and with appropriate restriction sites and cloned into PCR amplified plasmids with different RBS (rbsD, rbs31 or rbs33). Resulting constructs were named pAP33 through 38.

**Table 1.** Strains and plasmids used in this work

| Strain or Plasmid | Characteristic | Reference |
| --- | --- | --- |
| *S. meliloti* strains |  |  |
| Rm1021 | SU47 str-21 *expR102::IS*Rm2011-1 | (69) |
| Rm8530 | Rm1021 *expR*<sup>+</sup> | (16) |
| Rm11609 | Rm8530 *exoY::Tn5*-132 *wgaAB::Tn5* | (70) |
| RG26 | Rm8530 *exoY::pRG53* | This work |
| RG27 | Rm8530 Δ*exoY* | This work |
| RG33 | Rm8530 Δ*wgaAB* | This work |
| RG34 | Rm8530 Δ*exoY* Δ*wgaAB* | This work |
| RG35 | Rm8530 Δ*exoY* Δ*wgaAB* *rhaS::pRG73(wgaAB*<sup>+</sup>*)* | This work |
| Plasmids |  |  |
| pAP01 | P<sub>EL-222</sub>-*rbsD*-GFP in pSEVA531 backbone, Tc<sup>r</sup> | This work |
| pAP05 | pJ23105-*rbs34*-EI222 and P<sub>EL-222</sub>-*rbsD*-GFP in pSEVA531, Tc<sup>r</sup> | This work |
| pAP15 | pJ23115-*rbs34*-EI222 and P<sub>EL-222</sub>-*rbsD*-GFP in pSEVA531, Tc<sup>r</sup> | This work |
| pAP14 | pJ23105-*rbs34*-EI222 and P<sub>EL-222</sub>-*rbs34*-ATG-*wgaAB* in pSEVA531, Tc<sup>r</sup> | This work |
| pAP33 | placI<sup>q</sup>-*rbs34*-EI222 and P<sub>EL-222</sub>-*rbsD*-ATG-*wgaAB* pSEVA531, Tc<sup>r</sup> | This work |
| pAP34 | placI<sup>q</sup>-*rbs34*-EI222 and P<sub>EL-222</sub>-*rbs33*-ATG-*wgaAB* pSEVA531, Tc<sup>r</sup> | This work |
| pAP35 | placI<sup>q</sup>-*rbs34*-EI222 and P<sub>EL-222</sub>-*rbs31*-ATG-*wgaAB* pSEVA531, Tc<sup>r</sup> | This work |
| pAP36 | placI<sup>q</sup>-*rbsD*-EI222 and P<sub>EL-222</sub>-*rbsD*-GTG-*wgaAB* pSEVA531, Tc<sup>r</sup> | This work |
| pAP37 | placI<sup>q</sup>-*rbsD*-EI222 and P<sub>EL-222</sub>-*rbs33*-GTG-*wgaAB* pSEVA531, Tc<sup>r</sup> | This work |
| pAP38 | placI<sup>q</sup>-*rbsD*-EI222 and P<sub>EL-222</sub>-*rbs31*-GTG-*wgaAB* pSEVA531, Tc<sup>r</sup> | This work |
| pAP05EI222 | pJ23105-*rbs34*-EI222 in pSEVA531, Tc<sup>r</sup> | This work |
| pAP11GFP | pR-*rbs34*-GFP in pSEVA631, Gm<sup>r</sup> | This work |
| pRG52 | pGEM::Δ*exoY*, Ap<sup>r</sup> | This work |
| pRG53 | pJQ200SK::Δ*exoY*, Gm<sup>r</sup> | This work |
| pRG68 | pGEM::Δ*wgaAB*, Ap<sup>r</sup> | This work |
| pRG70 | pJQ200SK::Δ*wgaAB*, Gm<sup>r</sup> | This work |
| pRG73 | pCAP77::*wgaAB*<sup>+</sup>, Nm<sup>r</sup> | This work |

<sup>a</sup> Tc<sup>r</sup>, tetracycline resistance; Gm<sup>r</sup>, gentamycin resistance; Ap<sup>r</sup>, ampicillin resistance; Nm<sup>r</sup>, neomycin resistance.

### Measurement of promoter strength in *S. meliloti*

Promoters were tested using a transcriptional fusion to the reporter enzyme β-glucuronidase (GUS) (47), encoded by *gusA* (see **Supp. Table 1)**. The promoter-*gusA* constructs were generated using overlap extension PCR with primers containing the promoter of interest and *gusA* overlap sequence (see primers used in **Supp. Table 2**). In all of the constructs, the same ribosome binding site, BBa_B0034 (rbs34; parts.igem.org), was used. The resulting constructs were cloned into plasmid pSEVA531 (48) using Golden Gate (GG) assembly (49) (see list of resulting plasmids in **Supp. Table 2**). The pSEVA531 plasmid contained the pBBR1 origin of replication and *tetA* for selection. Freshly made *S. meliloti* RG34/pAP00GUS, RG34/pAPP_El222_GUS, RG34/pAP05GUS, RG34/pAP08GUS, RG34/pAP10GUS, RG34/pAP15GUS, RG34/pAPFixK2GUS, RG34/pAPlacI^q^GUS and RG34/pAPRGUS cells expressing *gusA* were inoculated into TY medium with tetracycline and grown for 48 h at 30 °C with 230 RPM shaking. Cells were diluted to an optical density (OD_600_) of 0.1 and grown until reaching OD_600_ 0.5-0.6, measured with a ND-1000 spectrophotometer with pathlength of 1 cm. Subsequently, cells were either used immediately for the GUS assay or stored at −80 °C in 0.5 mL aliquots. Frozen cells were thawed on ice, diluted to OD_600_ 0.3, and 500 μL of cell suspension was permeabilized by adding 30 μL 0.1 % SDS (w/v) and 60 μL chloroform and vortexing for 10 s. 50 μL of permeabilized cell suspension was added to 950 μL of GUS assay buffer (50 mM NaPO_4_ pH 7, 5 mM DTT, 1 mM EDTA, 1.25 mM 4-methylumbelliferyl β-D-glucuronide(MUG)) and incubated in a 37 °C water bath for 5 min. The reaction was halted by transferring 100 μL of sample to 900 μL 0.4 M Na_2_CO_3_. Formation of 4-methylumbelliferone (4-MU) was measured from 100 μL of stopped reaction mixture in cell culture treated optical bottom black polystyrene microplates (Thermo Scientific, cat #165305) in a plate reader (Biotek HT Synergy) by using fluorescence at 365/460 nm (excitation/emission). Experiments were carried out in an environment with minimal light to avoid decomposition of MUG.

### Light controlled sfGFP expression

*S. meliloti* strains RG34/pAP01 and RG34/pAP05 were grown in 5 mL TY medium with tetracycline selection for 48 hours at 30 °C with 230 RPM shaking without light exposure (culture tubes were wrapped with aluminum foil). Subsequently cells were diluted 100 fold in 500 μL M9 medium with appropriate antibiotics and grown under static conditions for 48 hours at 30°C a 6-well plate (Corning cat #3516) illuminated with 6 W/m^2^ blue light (Thorlabs 470 nm light-emitting diode (LED) filtered with a Chroma 480/40 bandpass filter). Due to the high light sensitivity of the EL222, light intensities well below those used for exciting fluorescent proteins are needed to induce light-driven transcriptional unit (50, 51). Cells were collected by centrifugation and washed in fresh M9 once and sfGFP expression levels were measured by flow cytometry using a BD Biosciences LSRFortessa X-20 Cell Analyzer (FITC channel). Individual signals from 30,000 cells were collected and data were analyzed by using FlowJo (vX.0.7).

For kinetic studies of sfGFP expression, *S. meliloti* strain RG34 with pAP01, pAP05 or pAP15 plasmids were grown in aluminum-wrapped tubes 5 mL TY media for 48 hours with appropriate antibiotics at 30 °C with 230 RPM shaking. Subsequently, cells were diluted in M9 to an OD_600_ of 0.1 and cultured in 24 well plates with 5 W/m^2^ blue light illumination at 30 °C without shaking. A duplicate plate was maintained in the dark. Cells were harvested at each time point and treated with chloramphenicol (100 μg/ml) to stop protein translation and kept at 4°C. Fluorescence levels were measured by flow cytometry as above.

### Autoaggregation assay

*S. meliloti* RG34/pAP14, RG34/pAP33, RG34/pAP34, RG34/pAP35, RG34/pAP36, RG34/pAP37 and RG34/pAP38 were grown in 5 mL TY with tetracycline selection for 48 hours without any light exposure at 30 °C with 230 RPM shaking (culture tubes were wrapped with aluminum foil). Culture optical density was measured (OD_600initial_) then tubes were incubated 24 hours without shaking at 4°C. The OD_600_ of the upper 0.2 mL of culture media was measured (OD_600final_). The autoaggregation percentage was calculated as: 100[1-(OD_600final_/OD_600initial_)]. Great care was taken not to disturb cultures during 4°C incubation and final optical density measurements.

### Anthrone Assay

To quantify the carbohydrate content in culture supernatants, cultures were transferred to a conical tube and centrifuged at 4000 RPM (3320 rcf) for 10 min. The supernatant was collected and kept at 4°C until use, no longer than 2 days. 0.2 % anthrone was prepared fresh by dissolving 0.1 g of anthrone (CAS: 90-44-8) in 50 mL of 95% sulfuric acid. 50 μL of supernatant was mixed with 150 μL of 0.2% anthrone in 1.5 mL Eppendorf tubes and incubated at 4°C for 10 min followed by 100°C for 20 min in an oven. Tubes were allowed to cool to room temperature and the A_620_ of the reaction mix was measured with a plate reader. Glucose equivalents were calculated from a glucose calibration curve.

### Light Controlled EPS II production

*S. meliloti* strains RG34/pAP14, RG34/pAP33, RG34/pAP34, RG34/pAP35, RG34/pAP36, RG34/pAP37 and RG34/pAP38 were grown in 5 mL TY medium for 48 hours with appropriate antibiotic selection at 30 °C with 230 RPM shaking without light exposure (culture tubes were wrapped with aluminum foil). Cells were diluted 100-fold in fresh TY medium with appropriate antibiotics and continuously exposed to 6 W/m^2^ blue light for 48 hours at 30°C. Duplicate control cultures were grown under the same conditions in the dark. Following growth, EPSII production was measured using the anthrone assay.

### Duty Cycle Experiments

Cells from 20 % glycerol stocks were grown in 5 mL TY media for 48 hours with appropriate antibiotic selection at 30 °C, with 230 RPM shaking, without light exposure (culture tubes were wrapped with aluminum foil). The cells were diluted to an OD_600_ of 0.1 in RDM media with antibiotics and cultured in 500 μL volume in cell culture treated 24 well polystyrene plates. Plates were illuminated from bottom with blue light at 2 W/m^2^ intensity with illumination programs described in the text for approximately 44 hours at 28 °C without shaking. A duplicate plate incubated in the absence of light was used as a control. An Arduino microprocessor was programmed accordingly for each duty cycle experiment.

### Confocal Microscopy

Cells from 20 % glycerol stock were inoculated in 5 mL TY media and grown for 48 h with appropriate antibiotics at 30 °C with 230 RPM shaking without light exposure (culture tubes were wrapped with aluminum foil). Cells were diluted to OD_600_ of 0.1 in RDM medium and 500 μL were cultured in 8-chamber glass slides (Lab-Tek II) for approximately 48 h at 28 °C. Samples were either illuminated with 1 W/m^2^ blue light (Thorlabs 470 nm LED filtered with a Chroma 480/40 bandpass filter) or kept in the dark throughout the incubation. Slides were imaged with a Nikon A1R inverted confocal microscope (488 nm excitation, 525/50 nm emission) with a 20x/0.45 S Plan Flour ELWD objective. Step size of 2.175 μm was used to generate Z-stack images. NIS-Elements software was used to operate the instrument and capture the images.

### Computation of biofilm properties from confocal microscopy data

COMSTAT 2.1 (Release date July 1,2015) was used to assess biofilm morphology according to the provider’s instruction (52, 53). **Nd2** image files were converted to **ome.tiff** files using the ImageJ Bio-Formats plugin. The ImageJ Comstat2 plugin was used to analyze image files in terms of *BioMass* (μm^3^/μm^2^), *Thickness Distribution* (μm), and *Surface Area* (μm^2^). During the calculations *Connected Volume Filtering* was selected to eliminate contributions from non-continuous layers of biofilm, as they can be freely floating and not be associated with structured biofilms that formed on the bottom of the glass chamber.

### Patterning structured biofilm formation

Cells from 20 % glycerol stock were inoculated in 5 mL TY medium and grown for 48 h with appropriate antibiotics at 30 °C, with 230 RPM shaking, without light exposure (culture tubes were wrapped with aluminum foil). Then cells were diluted to OD_600_ of 0.1 in RDM medium with appropriate antibiotics and cultured in 500 μL cell culture in a treated 24 well polystyrene plate illuminated with 1 W/m^2^ blue light for approximately 44 h at 28 °C without shaking. Half of each well was covered with electric tape and the other half was left open to illumination. After approximately 48 h a small area of the well covering both illuminated and un-illuminated regions was scanned with a fully motorized fluorescence microscope capable of imaging large samples (Keyence BZ-X800/BZ-X810). The individual captured images were reconstituted into a single large image using the manufacturer’s software.

## Results and Discussion

### Synthetic constructs for light-driven gene expression in *S. meliloti*

To enable spatial and temporal control of EPS production, we sought to implement light-activated gene expression in *S. meliloti* strain Rm8530 (16). Light-sensitive photoreceptor domains have been reported in *Rhizobium leguminosarum* (54, 55) and *Bradyrhizobium* (55), but not in *Sinorhizobium* (55). A BLAST search of the *S. meliloti* genome did not yield any genes encoding light oxygen voltage (LOV) domains, so we sought other means to control EPS production with light (**Supp. Table 3**). The LOV domain containing protein EL222 (56), from the gram-negative bacterium *E. litoralis* HTCC2594, is a blue-light sensing protein with a C-terminal helix-turn-helix (HTH) DNA-binding domain representative of LuxR-type transcriptional regulator (57). EL222 has been developed as a light modulated transcriptional regulator in *E. coli* (51). EL222 can act as either an activator or repressor depending whether its binding site is located upstream (activator) of, or within (repressor) a promoter sequence. Based on this information, we constructed a blue-light inducible gene expression system in *S. meliloti* strain Rm8530 (**Fig. 1a**). In order to generate a single-plasmid enabling light-triggered gene expression, a constitutive promoter driving expression of *EL222* was desired. We tested the activity of a number of well-characterized promoters (**Supp. Table 1**) by driving the expression of the reporter enzyme β-glucuronidase (GUS)(47) in *S. meliloti.* Robust GUS activity was detected from promoters BBa_J23100 and pR (**Supp. Fig. 3**). Promoters BBa_J233105, and pLacI^q^ drove moderate levels of expression, while BBa_J233108, BBa_J233110, and BBa_J233115 led to weak expression (**Supp. Fig. 3**). To assess light-activated gene expression in *S. meliloti*, we constructed a plasmid (pAP05, **Fig. 1b**), using the pSEVA531 backbone (58), which contained the pBBR1 origin of replication and *tetA* for selection. In pAP05, an EL222-binding promoter (P_El222_, **Supp. Table 1**) regulates the expression of superfolder GFP (sfGFP), while EL222 is constitutively expressed from the moderate-strength promoter BBa_J23105 (**Fig. 1b**, **Supp. Fig. 3**). The EPSI/II^-^ *S. meliloti* strain RG34, which makes no EPS, was transformed with pAP05, and showed a 5.4-fold induction of GFP fluorescence after 48 hrs of blue-light illumination (**Fig. 1c**). Compared to the untransformed *S. meliloti* strain RG34, RG34/pAP05 showed a small (20-fold), but significant, leaky sfGFP expression under dark conditions (**Fig. 1c**, *P* = 0.0063, unpaired two-tailed t-test comparing RG34 vs. RG34/pAP05 dark). Similar levels of sfGFP fluorescence were detected either under light or dark conditions, in strain RG34/pAP01 which lacked the EL222 gene contained in pAP05 (**Fig. 1c**, *P* = 0.034, unpaired two-tailed t-test comparing RG34 vs. RG34/pAP01), suggesting that the leaky expression originated from low levels of transcription from P_EL222_. Interestingly, this leaky expression driven by P_EL222_ was not detected in GUS assays (**Supp. Fig. 3**, P_EL222_), possibly due to the high intracellular stability of sfGFP (59). To assess if the light intensity used for activating EL222 affected the growth of *S. meliloti* RG34/pAP11GFP, we obtained growth curves with and without light stimulation (**Supp. Fig. 4**). Strain RG34 harboring pAP11GFP, which constitutively expresses sfGFP (**Supp. Table 2**), showed no difference in cell growth with and without blue light stimulation over 48 hrs (**Supp. Fig. 4**). We then characterized the time response of light-driven gene expression by measuring sfGFP fluorescence in *S. meliloti*. Strain RG34/pAP05 showed low sfGFP expression after 8 hours of illumination and a 4-fold induction after 24 h (**Fig. 1d**).

**Figure 1.**
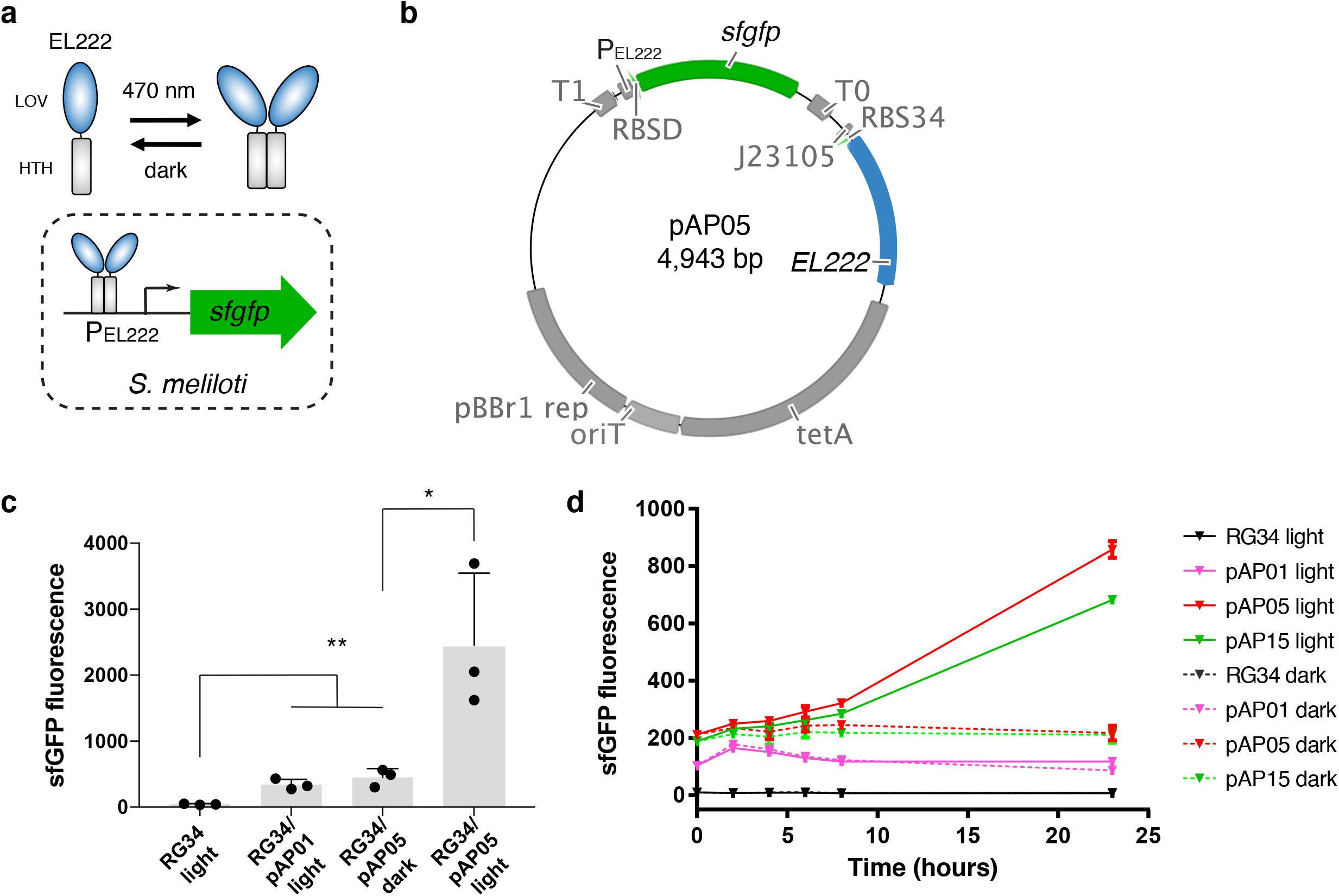
Light-driven gene expression in *S. meliloti.* **(a)** Schematic representation of blue light-driven transcription by EL222. EL222 consists of a N-terminal LOV domain (blue) connected to a C-terminal HTH DNA binding domain (grey) via a Jα helix (not shown). In the dark, EL222 is predominantly monomeric and unable to bind to DNA; however exposure to blue light shifts the equilibrium toward dimers, enhancing binding to the P_EL222_ promoter and transcription. **(b)** Schematic representation of plasmid pAP05 enabling blue light-driven expression of sfGFP. EL222 was expressed from the constitutive promoter BBa_J23105 and sfGFP from the P_EL222_ promoter. The plasmid backbone is based on pSEVA531. **(c)** Quantification of blue light-driven sfGFP expression in *S. meliloti* strains RG34, RG34/pAP01, and RG34/pAP05. As a control, RG34 was transformed with pAP01, which is identical to pAP05 except it is missing EL222. Cells were illuminated with blue light (6 W/m^2^) for 48 hours. Fluorescence intensities were then measured using flow cytometry. Statistics: *, *P* < 0.05; **, *P* < 0.01, unpaired two tailed t-test. **(d)** Time course of GFP levels of untransformed strains RG34, RG34 transformed with pAP01, pAP05 or pAP15. pAP15 is similar to pAP05 except EL222 expression is driven from weaker constitutive promoter (Supp. Fig 1. and Supp. Table 3). Cells were illuminated with blue light (solid line, 6 W/m^2^) or kept in dark (dashed line). Data plotted are mean ± standard deviation (n = 3 independently prepared samples).

### Optogenetic control of EPS production

To enable optical control of EPS II production, we sought to control the expression of *wgaAB* using the EL222 system in strain RG34. We determined the start codon of *wgaAB* based on a previous study that identified the gene cluster involved in EPS production and predicted the start codons of those genes (17). We assessed the potential leakiness of P_EL222_ when driving transcription of genes needed for EPS II synthesis as well its ability to activate light-driven EPS production by replacing *sfGFP* in pAP05 with the *wgaAB* genes (**Supp. Table 2**, **Fig. 2a**). Since we knew that a low, but significant, amount of leaky expression occurred when P_EL222_ drove *sfGFP,* we generated variants with different start codons (GTG and ATG) and inserted alternative RBSs in front of the *wgaAB genes* (**Fig. 2b**). To quantify EPS production, we used two previously developed assays (41, 60). First, we applied a sedimentation assay, which relies on aggregation of EPS-producing cells in planktonic culture (41). The sedimentation assay was effective in detecting leaky expression under dark conditions. We observed significantly higher sedimentation in strain RG34 transformed with plasmids pAP34, pAP35, pAP37, and pAP38, compared to the EPS deficient RG34 control cells (**Fig. 2c**), indicating that these plasmids had leaky production of EPS under dark conditions. The degree of sedimentation (as quantified by the sedimentation coefficient) showed a nearly binary response, where leaky constructs showed nearly complete sedimentation under dark conditions (RG34/pAP34, RG34/pAP35, RG34/pAP37, and RG34/pAP38), while sedimentation of RG34/pAP14, RG34/pAP33, and RG34/pAP36 was comparable to that of the EPS I and II-deficient parental strain RG34 (**Fig. 2c**). To compare the light-driven EPS production in strains that did not show significant leaky EPS production (RG34 with pAP14, pAP33, or pAP36), we measured the sugar concentration in culture supernatants using the anthrone assay (60). In this assay, significantly larger amounts of sugars were detected in strain RG27 (EPSI^-^, EPS II^+^) compared to the stain RG34 (EPS^-^, EPS II^-^) (**Fig. 2d**, *P* = 0.0044). Strain RG34/pAP14 exhibited a significant increase in culture carbohydrates upon blue light illumination (**Fig. 2d**, *P* = 0.037), indicating light-driven EPS production, while RG34/pAP33 and RG34/pAP36 did not show a significant increase (**Fig. 2d**, *P* = 0.26, *P* = 0.17, respectively). Strain RG34/pAP14 cells under dark conditions did not show a significant difference in sugar content compared to that of the EPS deficient RG34 cells (**Fig. 2d**, *P* = 0.36), indicating a lack of leaky EPS production. However, the amount of polysaccharide secreted by RG34/pAP14 cells under light was ~67% of that made by strain RG27 indicating modest EPS production recovery (**Fig. 2d**). Therefore, changing the start codon and the RBS of *wgaAB* did impact the ability to control EPS production, and as a result identified plasmid pAP14 (P_El222__*rbs34*_ATG_*wgaAB*) as having the best overall performance among those tested.

**Figure 2.**
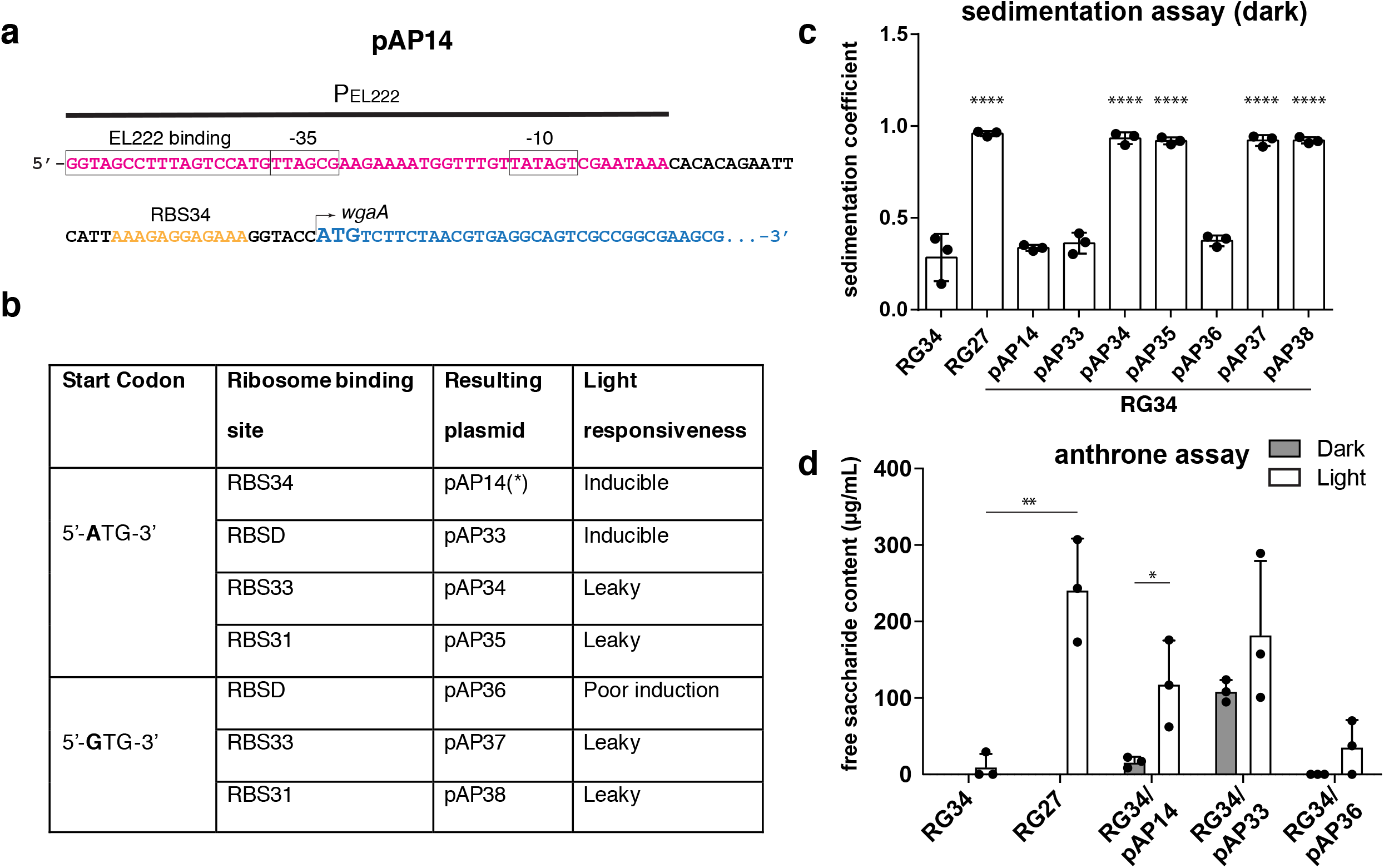
Quantitative assessment of light-driven EPS production in *S. meliloti.* **(a)** Parts of pAP14 for light-driven expression of *wgaAB.* The plasmid was constructed by replacing the *sfGFP* from pAP05 with *wgaAB.* The start codon of *wgaA* is shown in bold. **(b)** Plasmids generated to improve light-induced production of EPS. Plasmids that use an alternative start codon (GTG) as well as various ribosome binding sites (yellow sequence in (a)) are shown. The resulting plasmids were transformed into strain RG34 and assessed for light-induced production of EPS using the cell sedimentation assay. The ‘light responsiveness” column summarizes the results; ‘inducible’ indicates increased EPS production upon light stimulation, uninducible indicates no increase upon light activation, and ‘leaky’ indicates EPS production under dark conditions as inferred from the results in panel (c). **(c)** Sedimentation of planktonic *S. meliloti* cells in TY medium under dark conditions. **(d)** Measurement of secreted polysaccharide content using an anthrone assay. Free saccharide concentration was determined using glucose solutions as a standard. Plasmids pAP14, pAP33, and pAP36, which did not show leakiness in the sedimentation assay, were transformed in strain RG34, and tested for light-driven EPS production. Shaded bar graphs indicate samples kept in dark, while open bar graphs indicate samples continuously illuminated (470 nm LED, 6 W/m^2^). Plotted in panels (c) and (d) are mean +/− standard deviation of data from three independent culture preparations. Statistics for panels (c): ****: *P* < 0.0001 from ANOVA with Dunnett’s multiple comparisons test comparing the sedimentation coefficient of samples to that of strain RG34, (d): *: *P* < 0.05, **: *P* < 0.01, ns: *P* > 0.05 from unpaired t-test.

### Optogenetic control of mucoid phenotype and structured biofilm formation

As we expected, on solid media strain RG34/pAP14 showed a mucoid phenotype under blue-light illumination, but not under dark conditions, while strain RG34 exhibited a dry phenotype under both conditions (**Fig. 3a**). Another key characteristic of EPS-producing *S. meliloti* is the formation of three-dimensionally structured biofilms (30, 61). Previous studies have shown that EPS II is essential for biofilm formation and autoaggregation in *S. meliloti* (30, 41). Therefore, we assessed biofilm formation by the light-controlled production of EPS II. To visualize cells using fluorescence microscopy, we transformed a plasmid containing *sfGFP* under a constitutive promoter (pAP11GFP) into *S. meliloti* strains RG34 (EPS I^-^, EPS II^-^) and RG27 (EPS I^-^, EPS II^+^). The EPS II-producing strain RG27/pAP11GFP showed spatially heterogeneous biofilm formation, while strain RG34/pAP11GFP formed a uniform layer of cells (**Fig. 3b**, **Supp. Movies 1, 2**), consistent with previous observations (30). Strain RG34/pAP14+pAP11GFP showed similar spatial organization to the positive control strain RG27/pAP11GFP upon light activation (**Fig. 3b**, **Supp. Movie 3**). These results indicated successful light-driven control of EPS production in *S. meliloti* to an extent that allowed the formation of structured biofilms.

**Figure 3.**
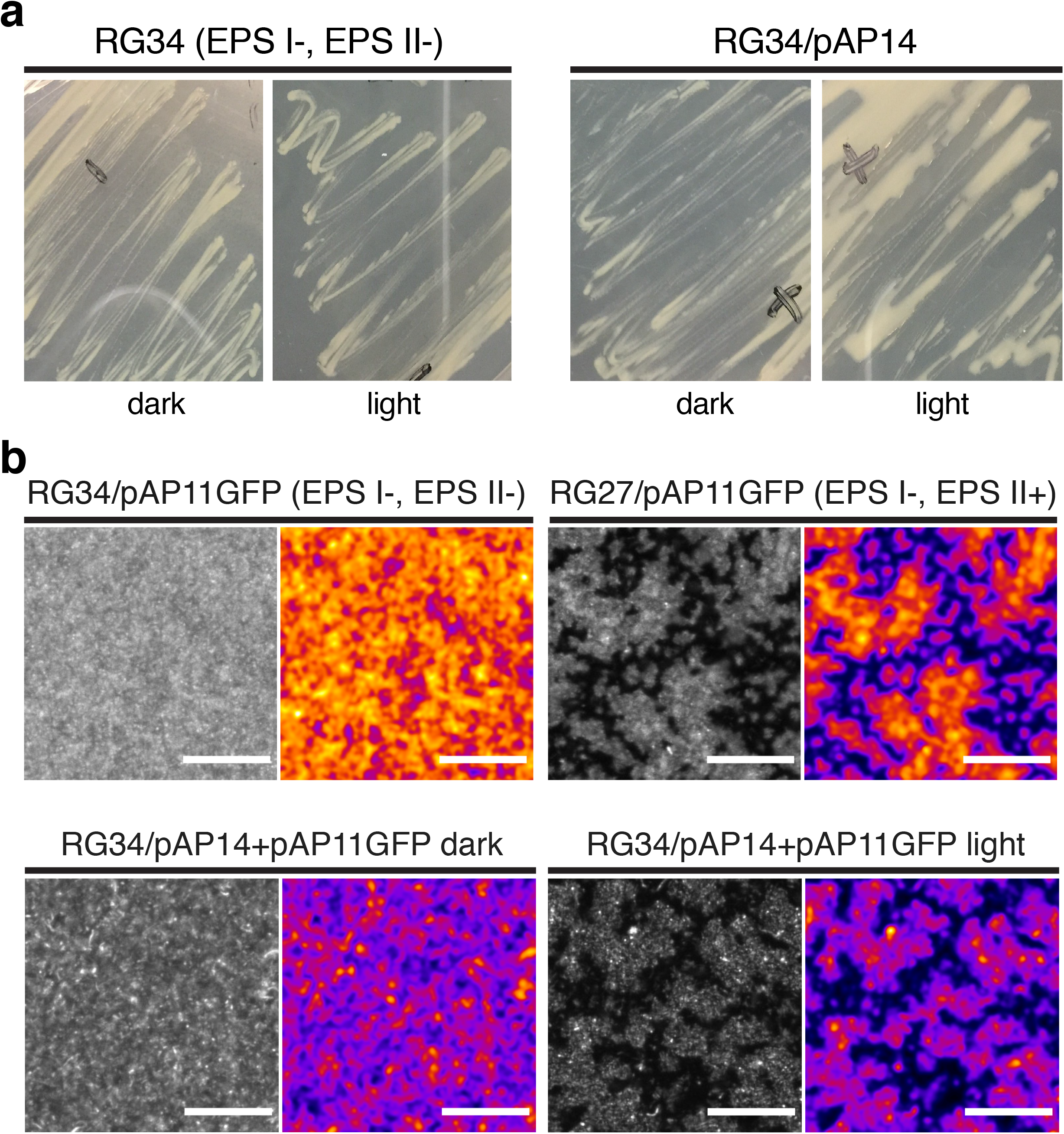
Optogenetic control of mucoid phenotype and biofilm structure in *S. meliloti.* **(a)** Assessment of EPS production on agar plates. Strains RG34 and RG34/pAP14 were grown on TY agar plates either under dark or with continuous illumination of blue light (470 nm LED, 6 W/m^2^) for 2 days. **(b)** Observation of biofilm structures by fluorescence microscopy. Strains RG34/pAP11GFP and RG27/pAP11GFP. Strains RG34/pAP14+pAP11GFP grown under dark conditions or with continuous illumination of blue light (470 nm LED, 6 W/m^2^). Fluorescence from sfGFP was imaged after 40 hrs. Scale bars in panel (b) indicate 20 μm. Grayscale images indicate sfGFP fluorescence and the pseudo-colored images are Gaussian filtered (ImageJ) images that enhance differences between biofilm structures. Time-lapse movies of these samples over 40 hrs are in **Supp. Movies 1-3** (Movies of RG34/pAP11GFP, RG27/pAP11GFP, RG34/pAP14+pAP11GFP, respectively).

Previous biophysical characterizations of the EL222 dynamics showed that the structural changes in EL222 upon light activation were spontaneously reversed in dark with a decay time constant (τ) of 11 s at 37°C (62). Consequently, transcriptional activity regulated by EL222 had a deactivation time constant of around 50 s (63). Based on these observations, we hypothesized that EPS II production and biofilm formation by *S. meliloti* could be controlled with pulses of blue light followed by dark periods, instead of continuous illumination. Although we have not observed blue-light mediated growth inhibition in *S. meliloti* at the light intensities used for activation of EL222, prolonged illumination may hinder long-term experiments due to phototoxic effects. Starting from continuous illumination (100% duty cycle), we reduced the duty cycle of illumination to 5% (3 s illumination followed by 57 s dark). Even with a 5% duty cycle, strain RG34/pAP14+pAP11GFP formed structured biofilms (**Fig. 4a**) that showed the same structured organization as biofilms made by strain RG27/ pAP11GFP producing EPS II (**Fig. 4b**). To quantify the structural organization, we measured the peak heights of the fluorescence profiles. Under dark conditions strain RG34/pAP14 + pAP11GFP showed low peak heights comparable to that of the EPS deficient RG34/pAP11GFP cells. The EPS II producing strains RG27/pAP11GFP and RG34/pAP14+pAP11GFP exposed to a 5% duty cycle of illumination showed significantly higher peak heights values compared to that of RG34/ pAP11GFP cells (**Fig. 4c**). These results suggested that short pulses of light with 5% duty cycle generated structured biofilms.

**Figure 4.**
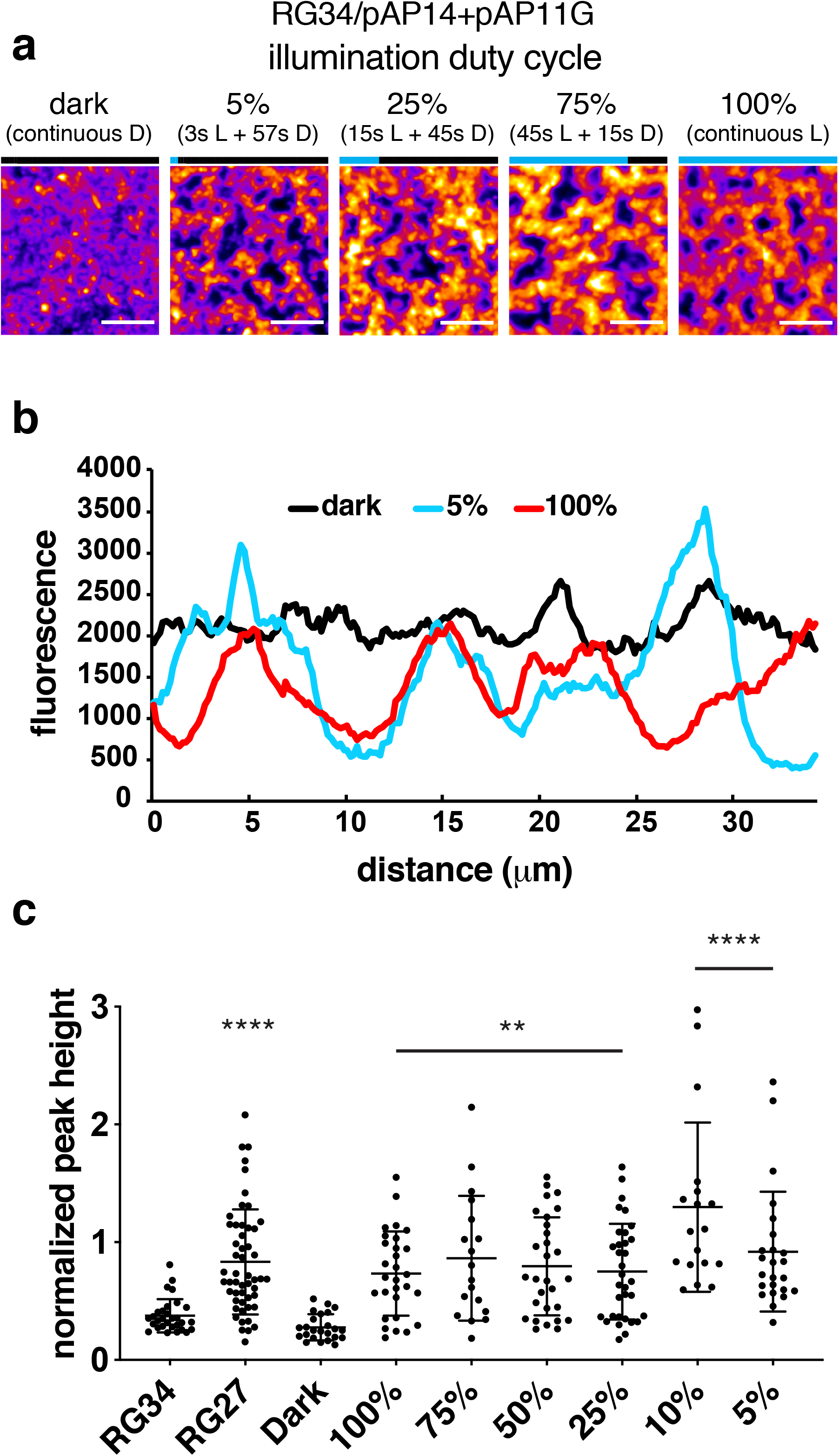
Structured biofilm formation with reduced illumination duty cycle. **(a)** Fluorescence images of biofilms under varying duty cycles. Duty cycles were calculated by the fraction of time cells were illuminated within each minute. 5% duty cycle indicates 3 s of light (L) followed by 57 s of dark (D). All cells were illuminated for a total duration of 48 hrs. Scale bars indicate 20 μm. **(b)** Fluorescence profiles quantified from images in panel (d). **(c)** Normalized peak heights quantified from fluorescence profiles. Data plotted are mean ± standard deviation of peaks quantified from n = 2-3 independently prepared samples. Statistics: **, *P* < 0.01; ****, *P* < 0.0001, Dunnett’s multiple comparisons test.

### Biofilm structure, thickness, and spatial control

Optical activation is highly suitable for *in situ* control of biological processes in complex environments. For example, optical control of EPS II production would allow spatial and temporal control of structured biofilm formation, which may enable testing the causal role of EPS II in the interaction between plant root nodulation and bacterial invasion (64–66). Using confocal microscopy, we characterized the thickness of biofilms and biomass formed by optically-controlled EPS II-producing *S. meliloti* using COMSTAT2 (52, 53). In order to assess the effect of light-driven activation of *wgaAB* on biofilm formation, we tested strain RG34/pAP14+pAP11GFP and control strain RG34/pAP05EL222+pAP11GFP. The latter strain lacked light-driven *wgaAB* genes and expressed EL222 constitutively. Notably, this control strain isogenic to the test strain, allowing the assessment of spurious activation of other genes by EL222, or by blue light alone, that may impact biofilm formation. After culturing RG34/pAP14+pAP11GFP and RG34/pAP05EL222+pAP11GFP for 2 days with and without blue light illumination, we observed cell clustering consistent with structured biofilm formation only in RG34/pAP14+pAP11GFP under blue light and high phosphate conditions (13 mM, **Fig. 5a**). Under low phosphate conditions (0.1 mM), which typically induce EPS II production, the structures within biofilm were not as clearly visible (**Fig. 5a**), perhaps due to overgrowth. Indeed, the low phosphate condition resulted in approximately 2-fold increase in biofilm thickness and biomass in all samples regardless of light exposure (**Fig. 5b, c**). Light activation induced a small, but significant increase in biofilm thickness (under both high and low phosphate conditions) and biomass (under high phosphate conditions) in RG34/pAP14+pAP11GFP but not in RG34/pAP05EL222+pAP11GFP cells (**Fig. 5b, c**). These results show that optical control of *wgaAB* expression resulted in biofilm properties consistent with previous studies using mutants that did or did not make EPS (30). Finally, we tested spatial control of structured biofilm formation by optically activating EPS production in defined regions of a sample. The EPS II inducible strain RG34/pAP14+pAP11GFP showed distinct structured biofilm formation only in illuminated regions, while the RG34/pAP05EL222+pAP11GFP control cells showed no difference in biofilm morphology in illuminated and dark regions (**Fig. 5d**). The transition from structured to non-structured regions occurred over approximately 400 μm (**Fig. 5d**), reflecting possible light scattering from the light region into the dark region, the diffuse nature of secreted EPS II and motility of *S. meliloti* cells during biofilm formation. Taken together, these experiments demonstrated spatial control of EPS II production and accompanying biofilm structure in *S. meliloti*.

**Figure 5.**
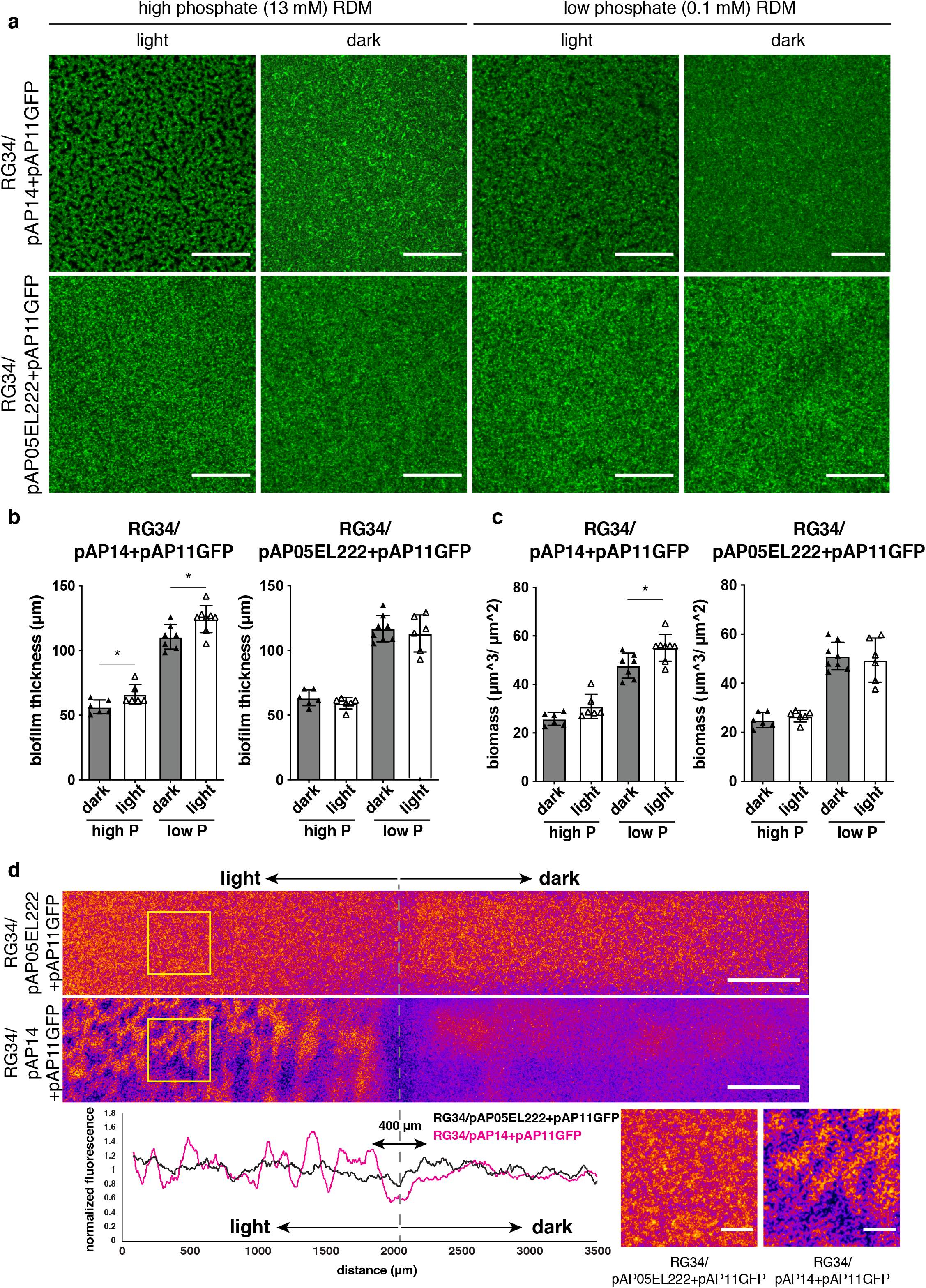
Spatial control of biofilm structure, thickness, and biomass in *S. meliloti*. **(a)** Confocal fluorescence images of light controllable biofilms formed under high- and low-phosphate conditions. Strains RG34/pAP14+pAP11GFP and RG34/pAP05EL222+pAP11GFP were grown in high or low phosphate RDM medium with or without light for 48 hours (470 LED 1 W/m^2^). The strain RG34/pAP05EL222+pAP11GFP (expressing EL222 only) was used as a negative control. Scale bars indicate 100 μm. **(b)** Biofilm thickness and biomass estimates, as measured by COMSTAT2, of biofilms shown in (a) **(c)** Spatial control of structured biofilm formation. RG34/pAP14+pAP11GFP and RG34/pAP05EL222+pAP11GFP were grown in high phosphate RDM for 44 hours. Black electrical tape was used to block light exposure of half of the 24 well plate (470 LED 1 W/m^2^). A region of the well covering both illuminated (light) and non-illuminated (dark) region of the plate was scanned. A fluorescence intensity profile of the region is shown. Scale bars in (d) indicate 500 μm (top panels) and 50 μm (bottom-right panels). Statistics for panel (b): *, *P* < 0.05, unpaired two tailed t-test.

## Conclusions

We have successfully implemented light-driven transcription in *S. meliloti* using the EL222 protein from *E. litoralis* and applied it to controlling the biosynthesis of EPS II. Optical control of EPS II synthesis was achieved by placing *wgaAB,* two genes required for its synthesis, under transcriptional control by EL222. Optical activation of EPS II production enabled spatial control of biofilm formation in *S. meliloti*, modifying biofilm thickness, biomass, and structure. EPS II biosynthesis was tightly controlled under dark conditions, and further refinement may enhance EPS II production upon light induction. The optical control approach shown here could be implemented for orthogonal gene expression control in *S. meliloti* when spatial and temporal regulation is desired. In particular, this approach can be easily implemented in recently developed experimental setups that allow optical access to plant root systems, soil, and bacteria (67, 68).

We anticipate that the tools developed here will lead to new insights on the role of EPS and other products in *S. meliloti* and related bacteria.

## Acknowledgements

This work was funded by the DOE grant DE-SC0014522. The authors have no conflicts of interest with the contents of this article.

## Supporting Information

### Supplementary Methods

#### Plasmid and strain construction

*Escherichia coli* XL1-Blue (Tet^r^) or XL1-Blue MRF’ (Kan^r^) (Stratagene) were used for all cloning steps. *E. coli* strains were routinely grown in LB medium (10g/L tryptone (Bacto), 5 g/L yeast extract (Bacto), 10 g/L NaCl (Fisher), solidified with 15 g/L agar (Bacto) when required), at 37°C. *Sinorhizobium meliloti* 8530 (1) (Rm8530 Str^r^) was used as the parental strain for construction of EPS deletion mutants and was routinely grown in TY medium (6 g/L tryptone, 3 g/L yeast extract, 0.38 g/L CaCl_2_ solidified with 15 g/L agar when required containing 500 μg/mL streptomycin sulfate (Sm)) at 30°C for 3-5 days. Broth cultures were shaken at 225 rpm (*E. coli*) or 150 rpm (*S. meliloti*). All strains were made electrocompetent as previously described (2). Plasmids were introduced by electroporation using approximately 10-100 ng plasmid DNA with an electric field strength of 1800 V (*E. coli*) or 2300 V (*S. meliloti*). Strains were allowed to recover in SOC medium (10 g/L tryptone, 2.5 g/L yeast extract, 0.6 g/L NaCl, 0.2 g/L KCl, 0.952 g/L MgCl2, 3.6 g/L D-glucose) for 1 hour (*E. coli*) or 4 hours (*S. meliloti*) at 37°C or 30°C, respectively. All kits were used according to manufacturer instructions. Plasmid pSEVA531 (3) was kindly provided by Victor de Lorenzo (National Center for Biotechnology (CNB), Madrid, Spain). Rm8530 genomic DNA (gDNA) was prepared (Promega Wizard Genomic DNA Purification Kit) and used as template for PCR. Deletions in mutant strains were confirmed by PCR and bi-directional dideoxy sequencing. Relevant strains are listed in the ‘Strains’ section of **Table 1**. Primers used are listed in **Supp. Table 2**.

#### Creation of the Rm8530 Δ*exoY* mutant

To create a mutant deficient in EPS I production an overlap extension protocol was used. The 5’ and 3’ ends of *exoY* (SMb20946) were amplified separately using Phusion DNA polymerase (NEB) and primers exoY_5’_F_XhoI and exoY_5’_R to generate a 650bp fragment containing sequence upstream of *exoY* and containing the first four codons of the open reading frame, and primers exoY_3’_F and exoY_3’_R_BamHI to generate a 500bp fragment containing the last 43 codons of *exoY* and extending into downstream *exoF* (SMb20945). Internal primers exoY_5’_R and exoY_3’_F 12 bp tails contained complementarity binding regions, allowing the use of PCR overlap extension (4) to scarlessly stitch amplicons together in-frame. First-round amplicons were purified and external primers exoY_5’_F_XhoI and exoY_3’_R_BamHI were used in the second-round PCR to create the chimeric *exoY* recombination locus. Second-round amplicons were gel-extracted, A-tailed and cloned into pGEM-T Easy. Ligations were introduced into *E. coli* XL1-Blue by electroporation as described and plasmid-containing clones selected on agar-solidified LB containing 50 μg/mL ampicillin (Ap,), 200 μM IPTG and 100 μM X-gal. Single, clone-positive (confirmed by PCR), colonies were used to inoculate LB cultures containing 50 μg/mL Ap. Cultures were used for freezer stocks and plasmid preparation. Plasmid containing the fused 5’ and 3’ *exoY* coding sequences was isolated and named pRG52. Plasmid pRG52 was double-digested with *BamHI-HF* and *XhoI* to excise the Δ*exoY* fragment and cloned into the suicide plasmid pJQ200SK (5) that had been digested in the same manner in the presence of rSAP. The Δ*exoY* fragment from pRG52 was ligated into the linearized pJQ200SK and the resulting plasmid was named pRG53. Plasmid pRG53 was introduced into Rm8530 by electroporation and cells were plated on TY supplemented with 30 μg/mL Gm to select for single-crossover mutants. 100 μL of sterile 10X M9 salts (58 g/L Na_2_HPO_4_ (Fisher), 30 g/L KH_2_PO_4_ (Fisher) 5 g/L NaCl, 10 g/L NH_4_Cl (Fisher)) was added to provide phosphate levels needed to help suppress EPS II production. Plates were incubated 3 days at 30°C and single colonies screened for single-crossover events by PCR targeting pRG53 plasmid sequence using primers M13_F and exoY_3’_R_BamHI. Rm8530 containing integrated pRG53 was named strain RG26. To counterselect against plasmid pRG53, RG26 was grown in TY broth to stationary phase, serially diluted and plated on TY containing 5% sucrose. Sucrose-resistant colonies were screened by PCR using primers exoY_5’_F_XhoI and exoY_3’_R_BamHI and transformants that produced truncated *exoY* amplicons were purified on TY. Rm8530 containing an in-frame deletion of *exoY* from codons 5-184 was confirmed by dideoxy sequencing using primers exoY_seq_F and exoY_seq_R as described above and named strain RG27(Δ*exoY*).

#### Creation of Δ*wgaAB* mutant

Expression of the *wga* (formerly *expA*) operon has been shown to be required for EPS II production (1). To create a mutant deficient in EPS II production, we targeted *wgaB* (SMb21320, formerly *expA*23), which encodes a putative glycosyltransferase. Due to the translational coupling of *wgaB* with the immediately upstream gene *wgaA* (SMb21319), the entire *wgaAB* locus including the *wgaAB* ribosome binding site (RBS) was targeted for deletion to reduce the risk of inefficient expression of *wgaB* from the *wga* promoter during complementation of the deletion. Rm8530 gDNA template was amplified using Phusion polymerase and the primer sets wgaAB_5’_F_SpeI and wgaAB_5’_R which amplify a 1.1 kb fragment containing the 5’ end of *wgcA* and most of the *wga* promoter, or wgaAB_3’_F and wgaAB_3’_R_NotI to amplify a 1.0 kb fragment containing most of the downstream gene *wgaD* (SMb21321). Primers wgaAB_5’_R and wgaAB_3’F were designed with 19 bp complimentary overlapping sequence which reconstitutes the *wgaD* RBS such that the final Δ*wgaAB* mutant expresses *wgaD* and remaining downstream *wga* genes directly from the *wga* promoter. The 5’ and 3’ fragments of *wgaAB* were gel purified and stitched together using external primers wgaAB_5’_F_SpeI and wgaAB_3’_R_NotI and cloned into pGEM-T Easy. Ligations were introduced into *E. coli* XL1-Blue and transformants screened for plasmid as described above. Transformants were screened and purified as described above and the plasmid named pRG68. pRG68 was double-digested using *NotI-HF* and *SpeI*-HF (NEB) to excise the Δ*wgaAB* fragment. pJQ200SK was linearized using the same enzymes in the presence of rSAP. The Δ*wgaAB* fragments were ligated to linearized pJQ200SK and introduced into *E. coli* XL1-Blue as described above. Transformants were screened and purified and the plasmid named pRG70. Single and double crossovers of pRG70 in *S. meliloti* strain Rm8530 were made using the same methods described above. Strain Rm8530 containing a complete deletion of *wgaAB* open reading frames was confirmed by PCR and dideoxy sequencing using primers wgcA_seq_F and wgaD_seq_R as described above and named strain RG33.

#### Creation of Δ*exoY*Δ*wgaAB* double deletion mutant

To create a mutant deficient in both EPS I and EPS II biosynthesis, *S. meliloti* ΦN3 (6) was used to transduce the single crossover *exoY*::pRG53 in *S. meliloti* strain RG26 into strain RG33 (Rm8530 Δ*wgaAB*). Strain RG26 was grown to late exponential phase in TY broth containing 30 μg/mL Gm at 30°C. Serial 10-fold dilutions of ΦN3 in 10 mM MgSO_4_ were created and each dilution combined with 100 μL RG26 culture. Phage-cell mixtures were allowed to adsorb for 30 minutes at 30°C, and each was transferred to 2 mL melted TY soft agar (0.75% agar) and poured over LB plates containing 30 μg/mL Gm. Plates were incubated at 30°C until plaques were visible, and plates containing confluent lysis were flooded with 4 mL TY broth incubated at 4°C to elute phage particles containing RG26 genomic loci. Lysate was collected, passed through a 0.45 μm nylon filter and a few drops of chloroform (Sigma) were added to give a RG26 ΦN3 lysate. Strain RG33 was grown to late exponential phase at 30°C in TY broth and 200 μL volumes mixed with 0, 10, 50, 100 or 200 μL volumes of RG26 ΦN3 lysate. Phage were allowed to adsorb for 20 minutes at 30°C and 1 mL LB broth added to each tube. Cells were washed 3 times in LB broth and suspended in 75 μL of LB containing 10mM sodium citrate. Prepared cells were plated on TY without CaCl_2_ containing 30 μg/mL Gm and incubated at 30°C. Transductants were screened for the presence of *exoY*::pRG53 by PCR using GoTaq polymerase and primers M13_F and exoY_3’_R_BamI. Insert-positive colonies were purified on TY containing 30 μg/mL Gm. To allow plasmid excision and formation of Δ*exoY*, broth cultures of *RG33(ΔwgaAB, exoY*::pRG53) underwent sucrose counterselection as described above. Colonies were PCR screened for the *exoY* deletion using GoTaq and primers exoY_5’_F_XhoI and exoY_3’_R_BamHI and recombinants exhibiting a deletion in *exoY* were purified on TY. A single recombinant was selected and di-deoxy sequenced to confirm genotype Δ*exoY*Δ*wgaAB*. This *S. meliloti* EPSI and EPS II double mutant was named strain RG34.

#### Complementation of Δ*wgaAB* mutant

To ensure that RG33 (Δ*wgaAB*) and its derivatives could be complemented to restore the ability to biosynthesize EPS II, *wgaAB* was cloned onto a suicide plasmid and inserted into the *rhaS* (SMc02324) locus on the RG34 chromosome by single recombination (2). Briefly, Rm8530 gDNA was used as template to PCR amplify *wgaAB* with the native *wga* promoter region using Phusion polymerase and primers wgcA_comp_F and wgaD_comp_R, which contain 16 or 18 bp overhangs with complementarity to pCAP77, respectively. The 4.5 kb *wgaAB* amplicon along with *EcoRI/HindIII* digested pCAP77[2] were checked for correct size by gel electrophoresis and the remainder of the PCR and restriction digest column purified and assembled using Gibson Assembly. Assemblies were introduced into XL1-Blue MRF’ and transformants were screened using primers M13_R and wgaA_scrn_R. Plasmid was isolated from a single transformant exhibiting the correct amplicon and named pRG73. RG34 (Rm8530 Δ*exoY*Δ*wgaAB*) electrocompetent cells were prepared and transformed with pRG73. Recovered cells were plated on TY containing 100 μg/mL neomycin sulfate and incubated at 30°C. Several mucoid colonies resembling wild-type Rm8530 were streaked to singles on the same medium and M9 medium (1X M9 salts, 0.1 mM CaCl_2_, 1 mM MgSO_4_, 5 ng/mL CoCl_2_, 0.5 μg/mL biotin) containing 0.4% L-rhamnose, 500 μg/mL Sm and 100 μg/mL Nm. Transformants were screened for their inability to grow on rhamnose as the sole source of carbon. Isolates exhibiting robust growth on TY but poor growth on rhamnose were retained for PCR screening of the *rhaS* locus using GoTaq polymerase and primers rhaS_Nterm_F and wgaA_scrn_R. Candidates exhibiting the expected amplicon size were grown to stationary phase in LB broth containing 2.5 mM MgSO_4_, 2.5 mM CaCl_2_ and 100 μg/mL Nm. A single candidate exhibiting poor growth on rhamnose and the functional rescue of EPS II biosynthesis was selected and named strain RG35.

#### Confirmation of EPS I/II deficiency in deletion strains

To compare the succinoglycan (EPS I) biosynthesis phenotypes of parental and deletion strains, Rm1021 (7), Rm8530, Rm11609 (8), RG27 and RG34 were streaked on MGS medium (50 mM morpholinepropanesuflonic acid (MOPS) 19 mM sodium glutamate, 55 mM D-mannitol, 1 mM MgSO_4_, 0.25 mM CaCl_2_, 0.1 mM KH_2_PO_4_, 0.1 mM K_2_HPO_4_) containing 5 ng/mL CoCl_2_, 0.5 μg/mL biotin and 0.02% w/v Calcofluor white M2R (Sigma) and incubated for 5 days at 30°C. Photographs were taken in ambient light or under long-wave UV transillumination to observe fluorescence due to EPS I production (**Supp. Fig. 1**). To assess differences in the mucoid phenotype (EPS II) strains Rm8530, Rm11609, RG34 and RG35 were streaked on the same medium as for EPS I and allowed to grow 5 days at 30°C. Photographs were taken and dense growth was visually assessed for the typical mucoid phenotype resulting from EPS II biosynthesis (**Supp. Fig. 2**).

### Supplementary Figures

**Supplementary Figure 1.**
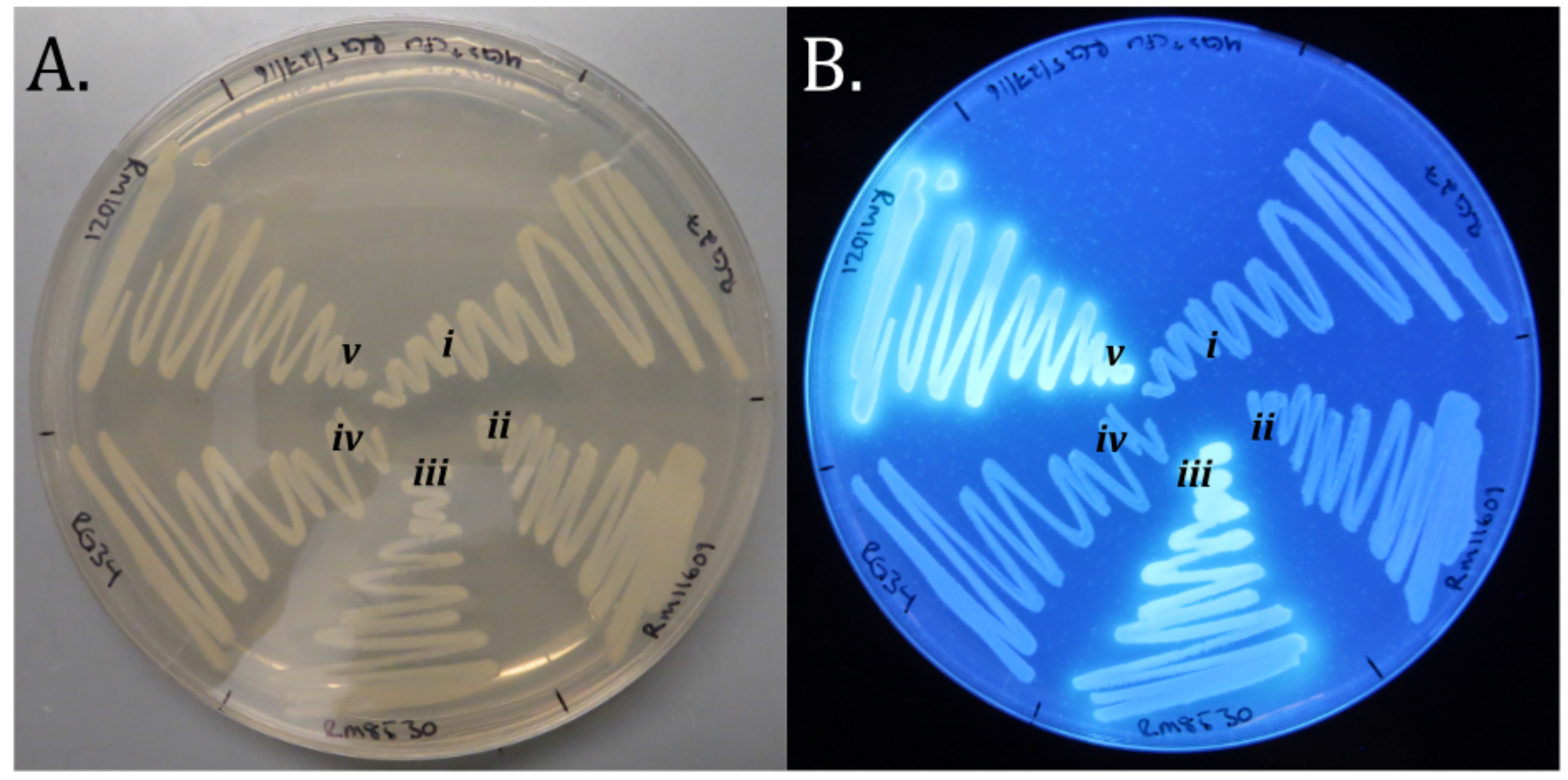
Δ*exoY* mutants lack succinoglycan (EPS I) biosynthesis. *S. meliloti* strains RG27(Δ*exoY*) (*i*), Rm11609(R*m8530 exoY::Tn5-132 wgaAB::Tn5*) (*ii*), Rm8530(wt) (*iii*), RG34(Δ*exoY* Δ*wgaAB*) (*iv*) and Rm1021(EPS II deficient) (*v*) were grown on MGS medium containing calcofluor white, which fluoresces under UV irradiation in the presence of succinoglycan. Photographs were taken under ambient (A) or long-wave UV transillumination (B). Insertional inactivation (*ii*) or in-frame deletion of *exoY (i, iv*) resulted in an EPS I deficient phenotype.

**Supplementary Figure 2.**
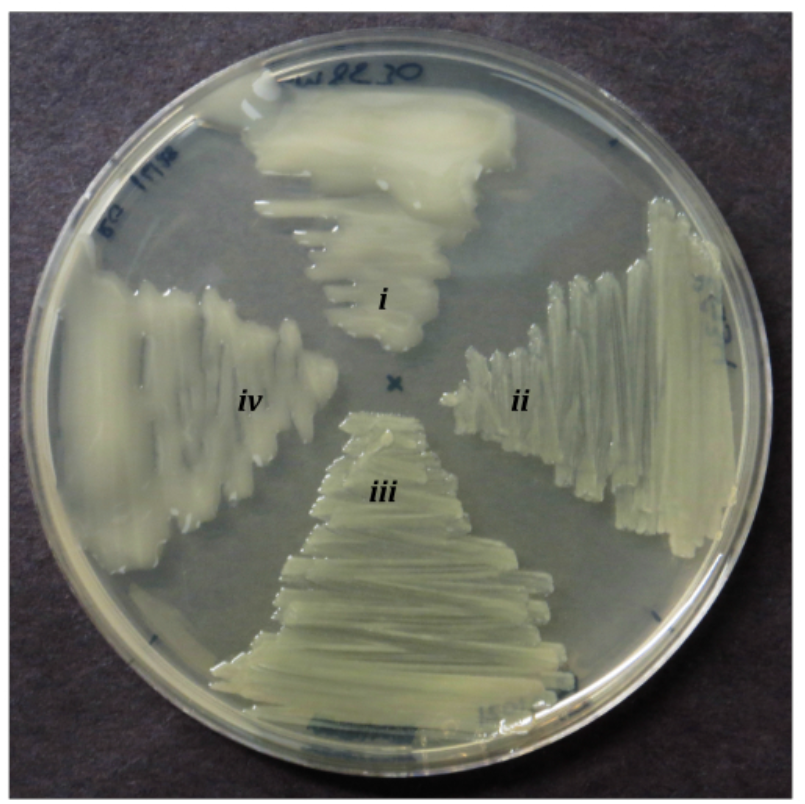
Complementation of Δ*wgaAB* from the *rhaS* locus rescues EPS II biosynthesis. *S. meliloti* strains Rm8530(wt) (*i*), RG34(Δ*exoY* Δ*wgaAB* (*ii*), Rm11609(R*m8530 exoY::Tn5-132 wgaAB::Tn5*) (*iii*) and RG35Δ*exoY ΔwgaAB rhaS*::pRG73(*wgaAB*^+^) (*iv*) were streaked on TY to observe mucoid phenotypes of each strain. Deletion of *wgaAB* is sufficient to inhibit the production of EPS II (*ii*). Rescue of EPS II biosynthesis is achieved through complementation of *wgaAB* under their native promoter and RBS by insertion into the *rhaS* (SMc02324) locus (*iv*).

**Supplementary Figure 3.**
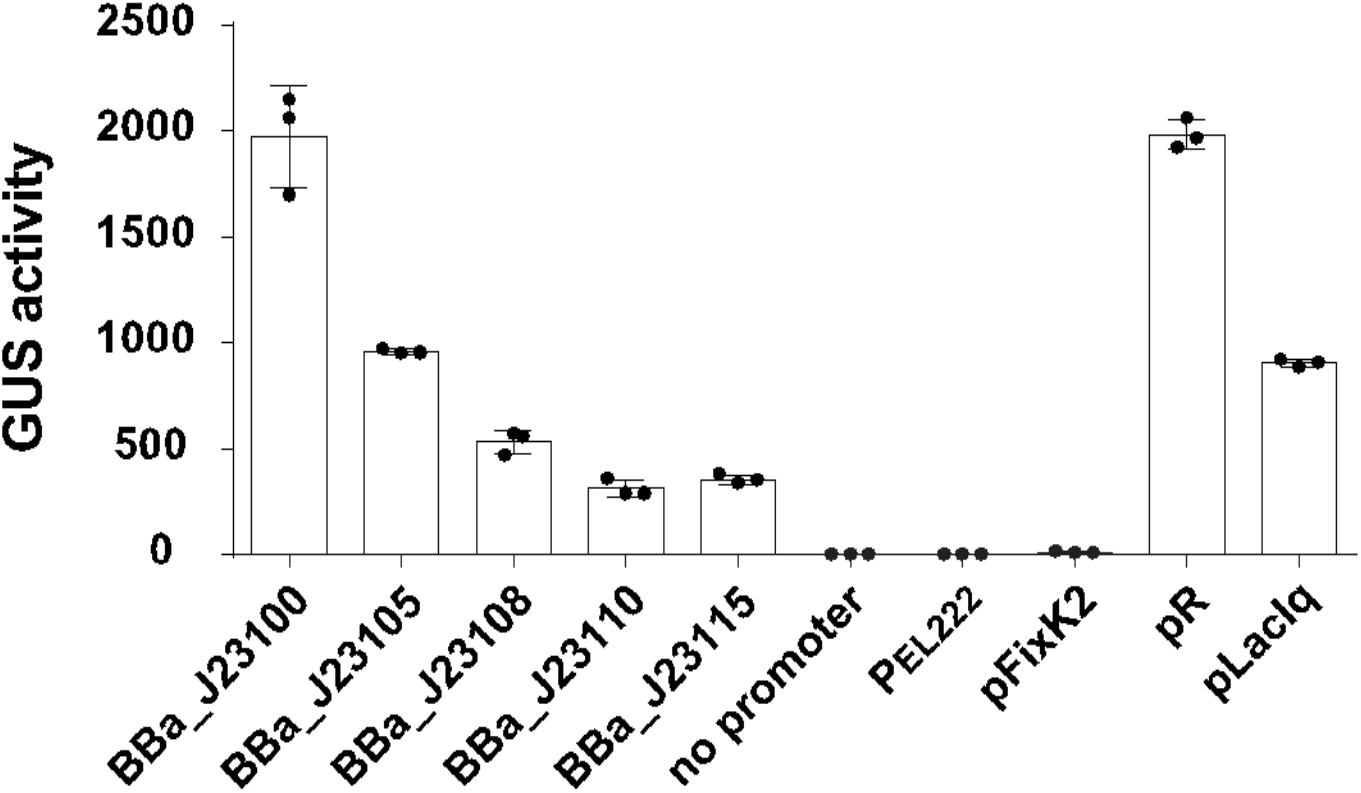
Assessment of promoter strength in *S. meliloti* using a β-glucuronidase (GUS) assay. *S. meliloti* strain RG34 was transformed with plasmids containing the indicated promoter driving constitutive expression of GUS. Data plotted are mean ± standard deviation (n = 3 independent culture preparations).

**Supplementary Figure 4.**
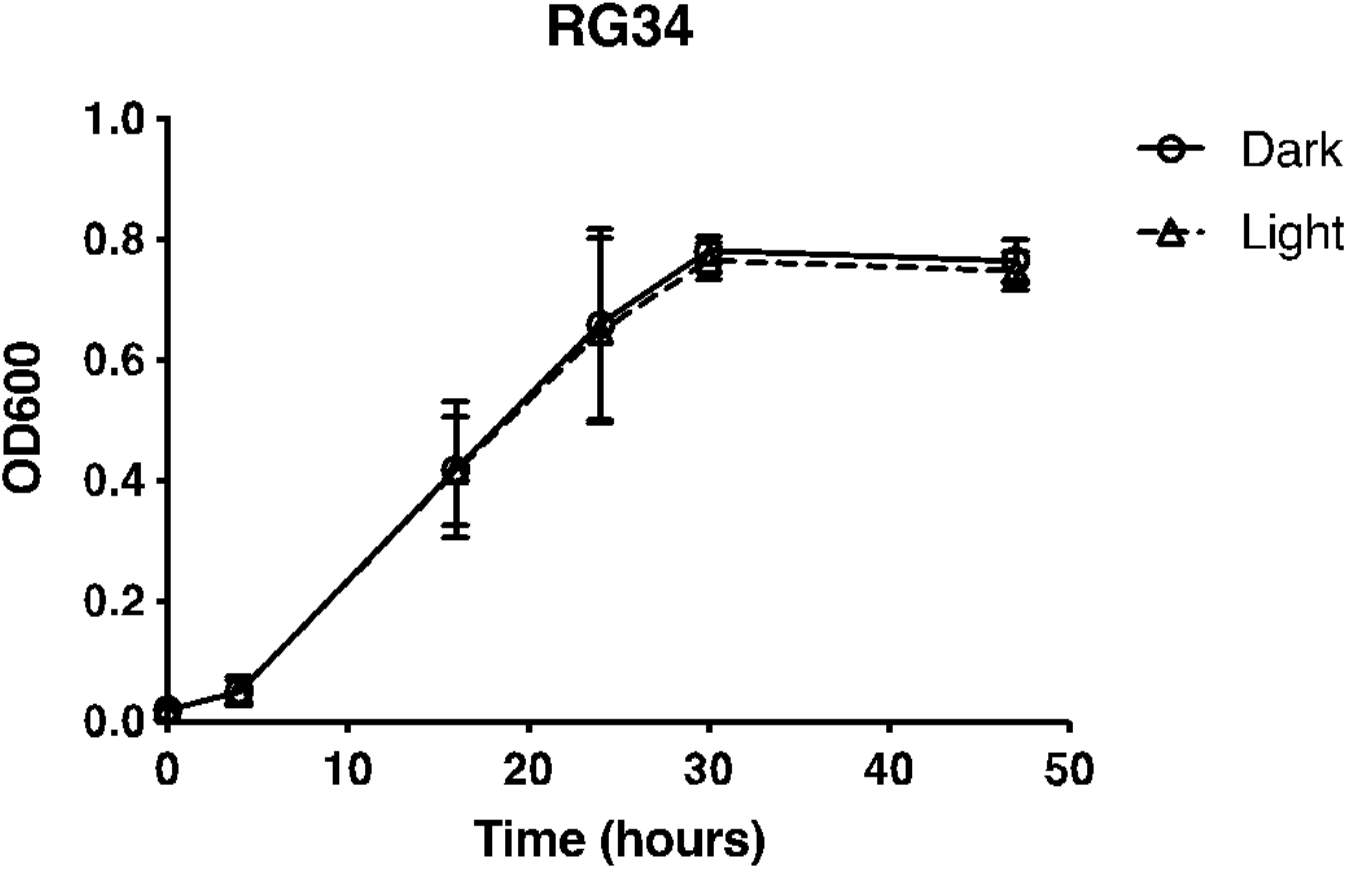
Comparison of *S. meliloti* growth with and without light stimulation. RG34 transformed with a sfGFP expressing plasmid under Gentamycin selection was grown over 48 hrs under constant blue light illumination (LED 6 W/m^2^). Cell growth quantified by absorbance at 600 nm (OD_600_) showed no difference (*P* = 0.830, repeated measures ANOVA, F(1,4) = 0.0528). Data plotted are mean ± standard deviation (n = 3 independent culture preparations).

### Supplementary Tables

**Supplementary Table 1.**
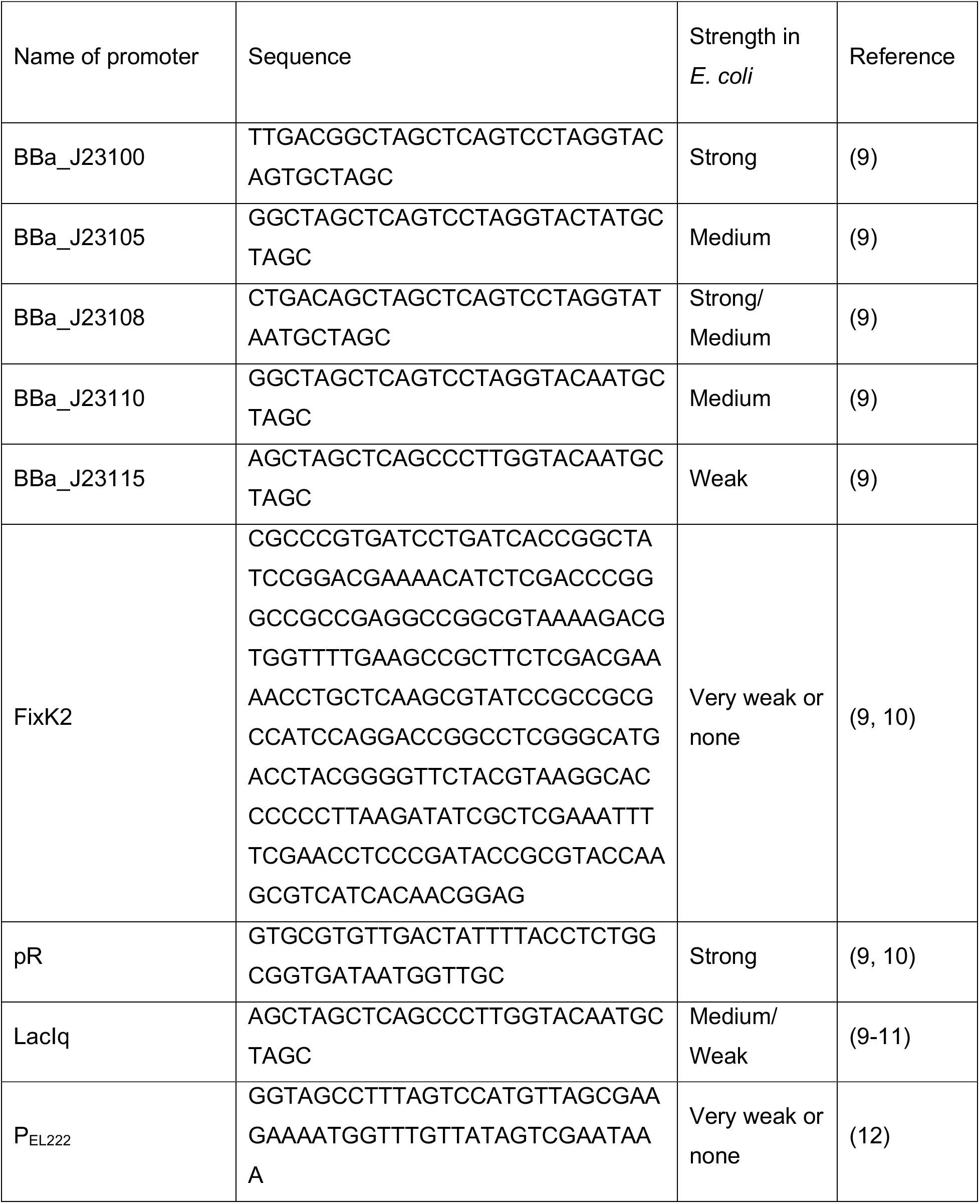
Promoters tested in *S. meliloti.*

**Supplementary Table 2.**
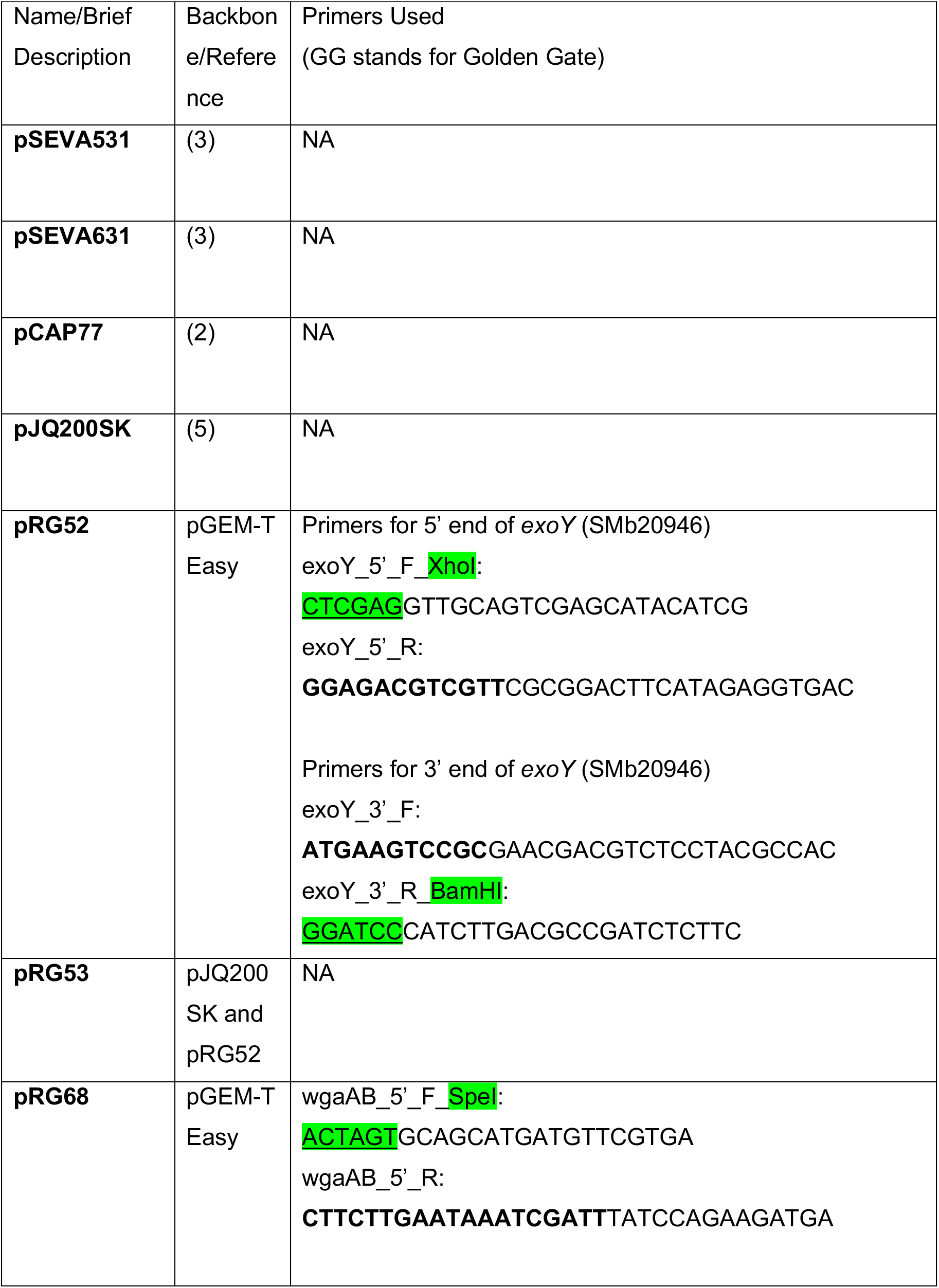
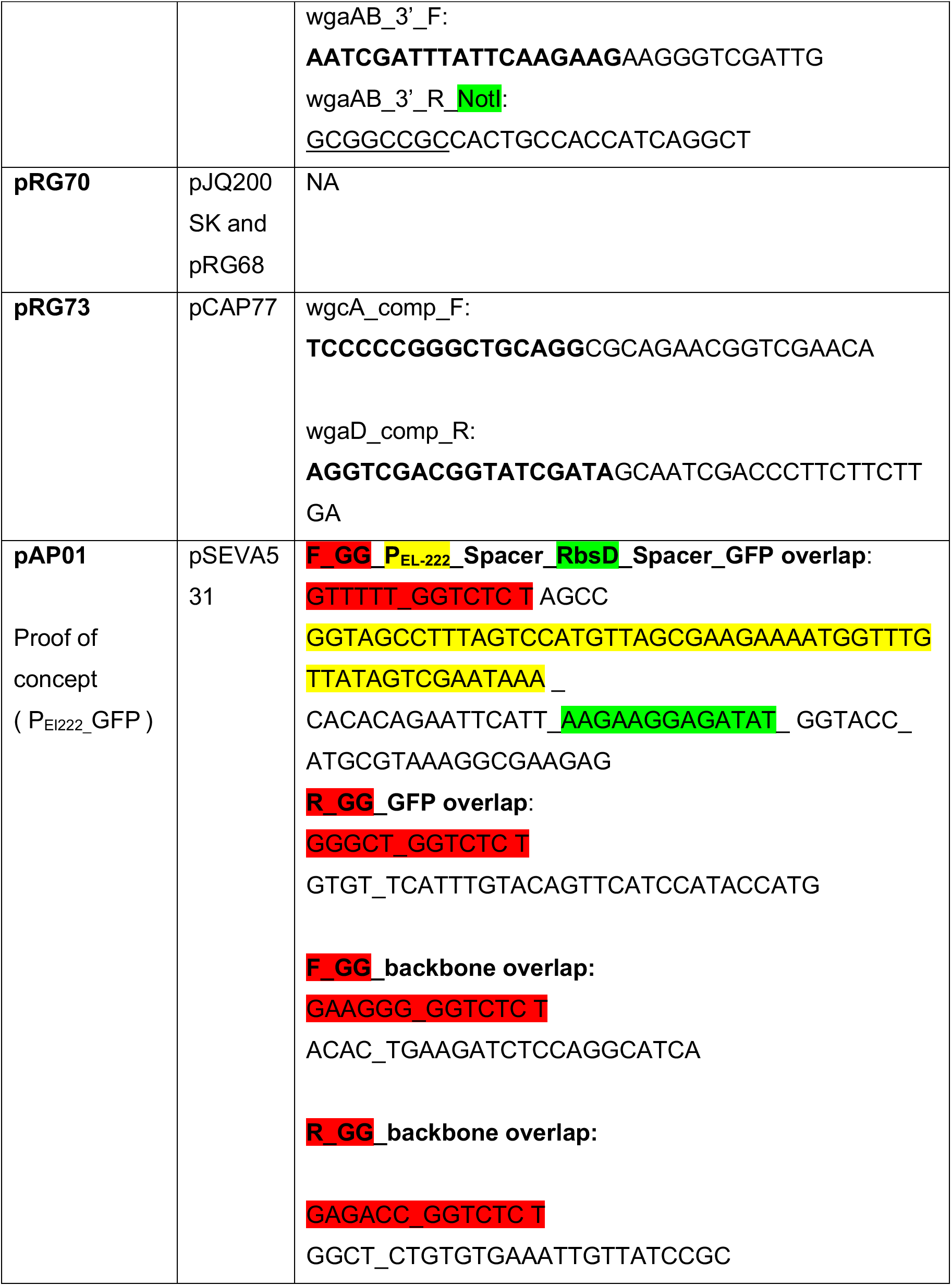
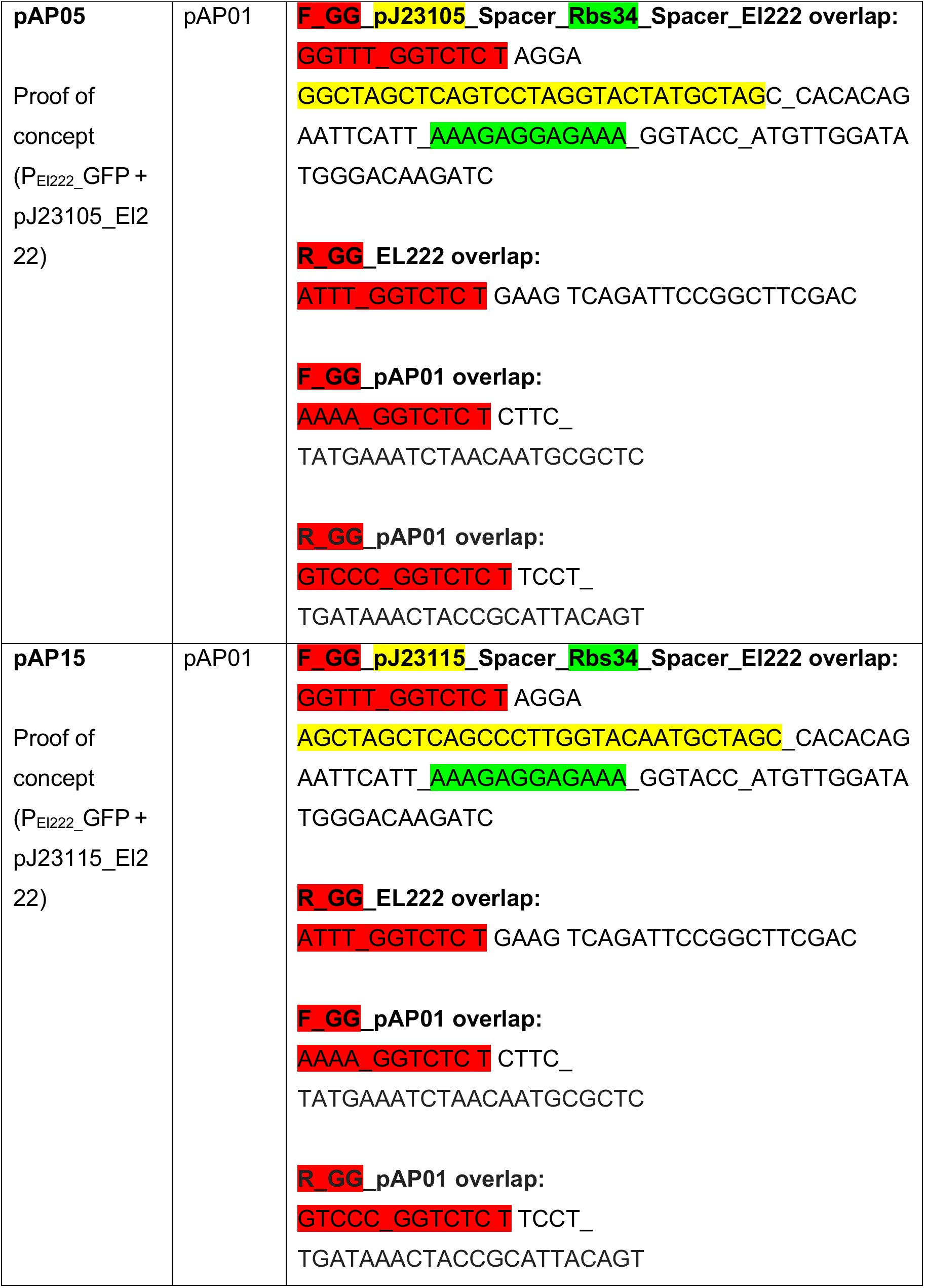
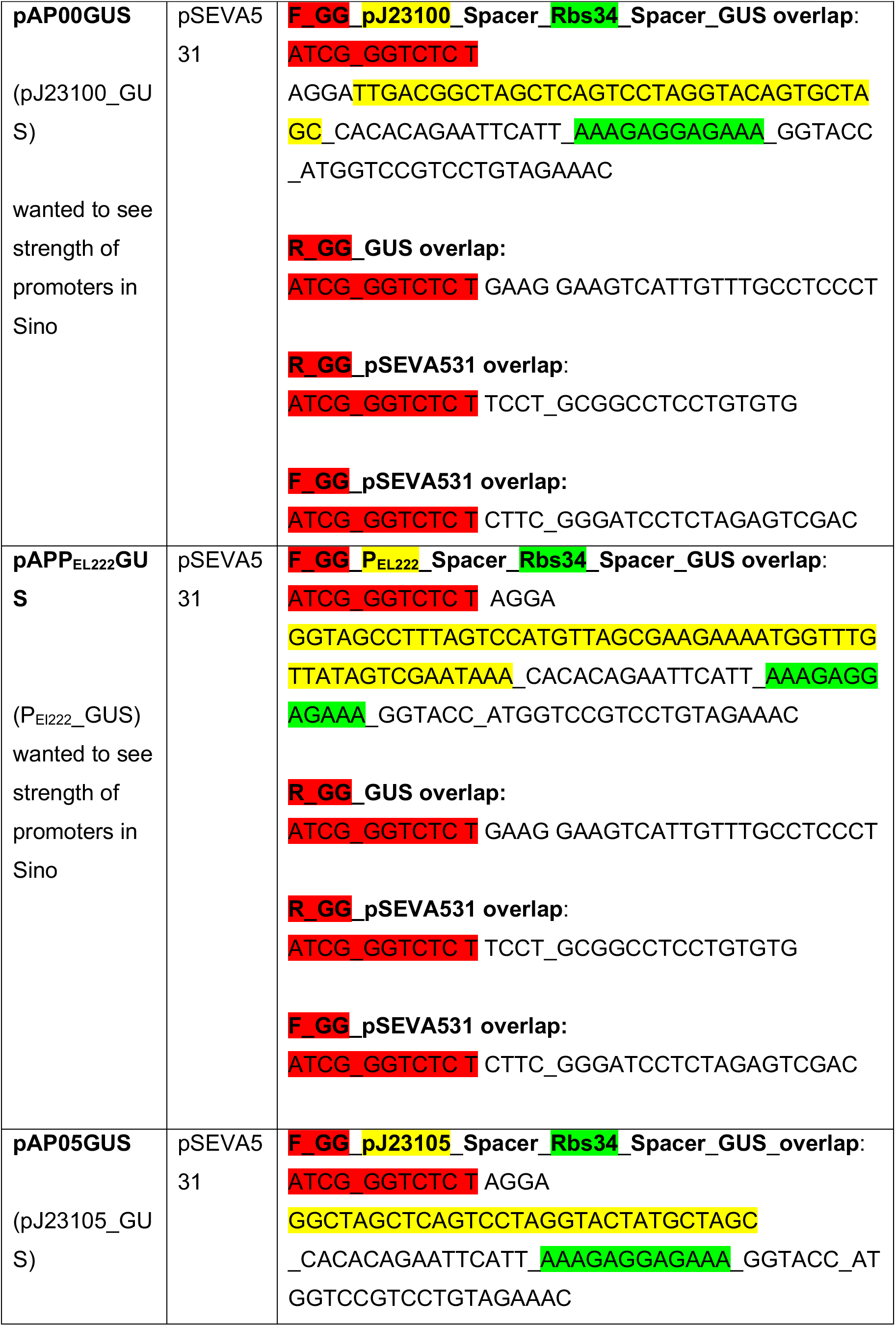
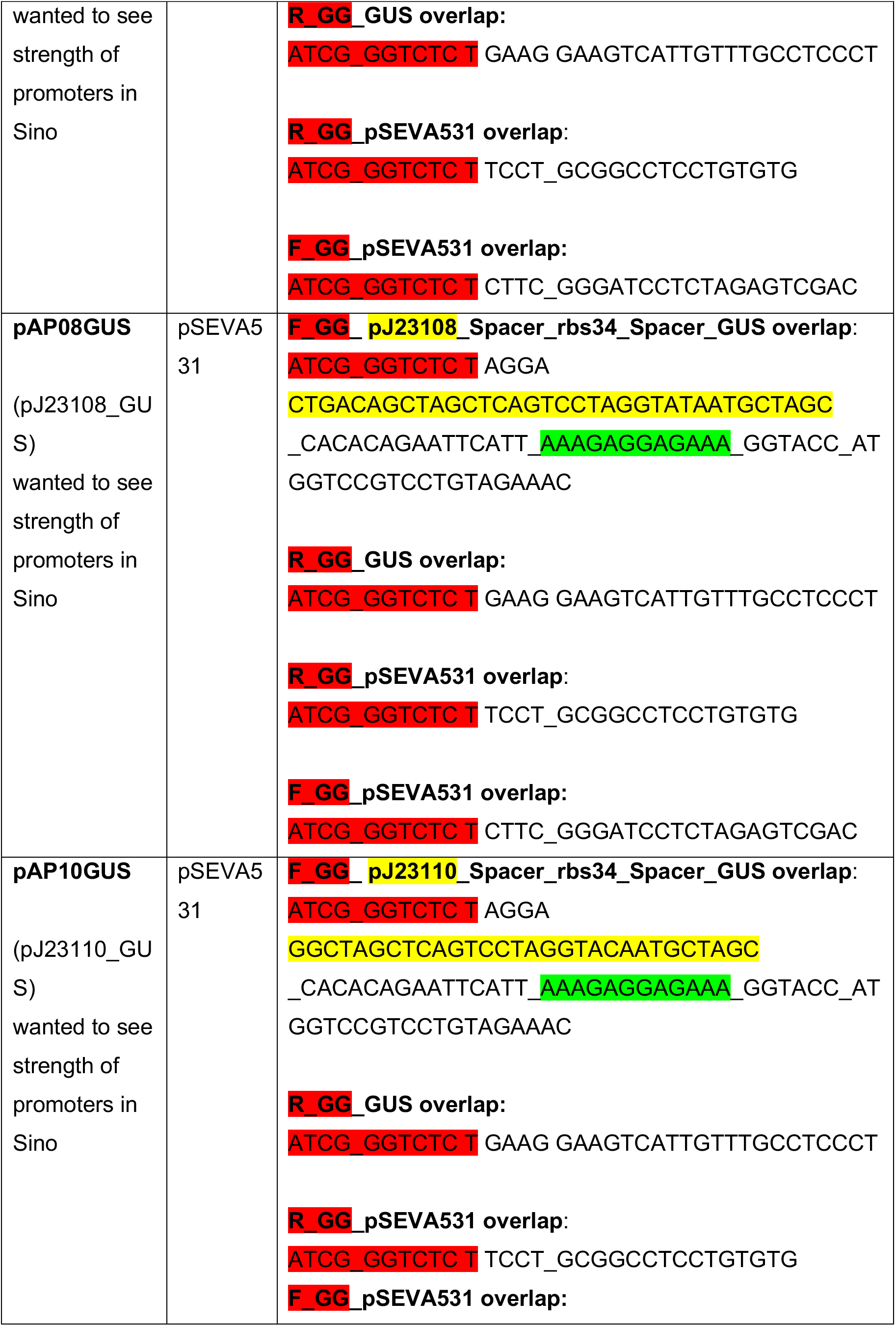
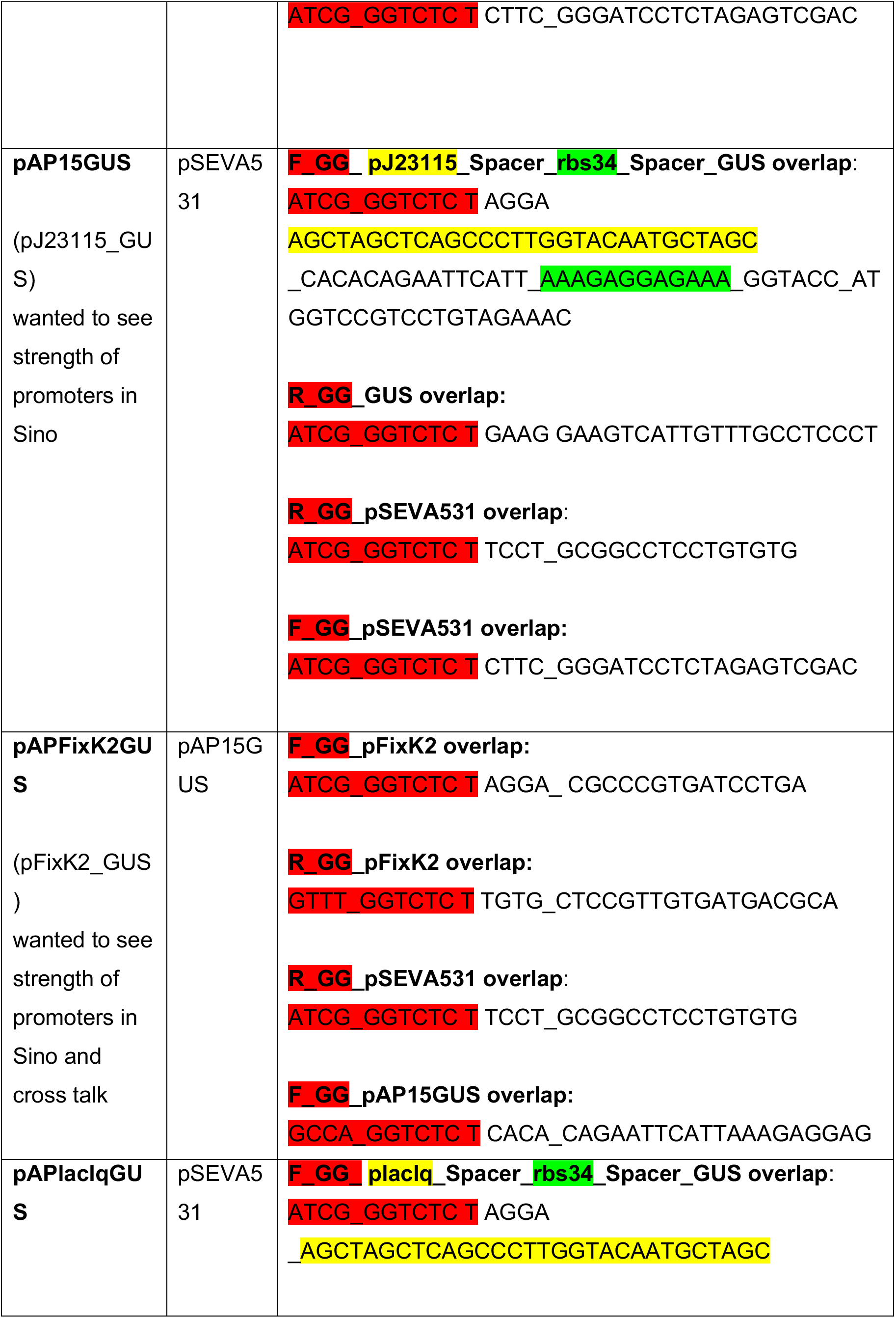
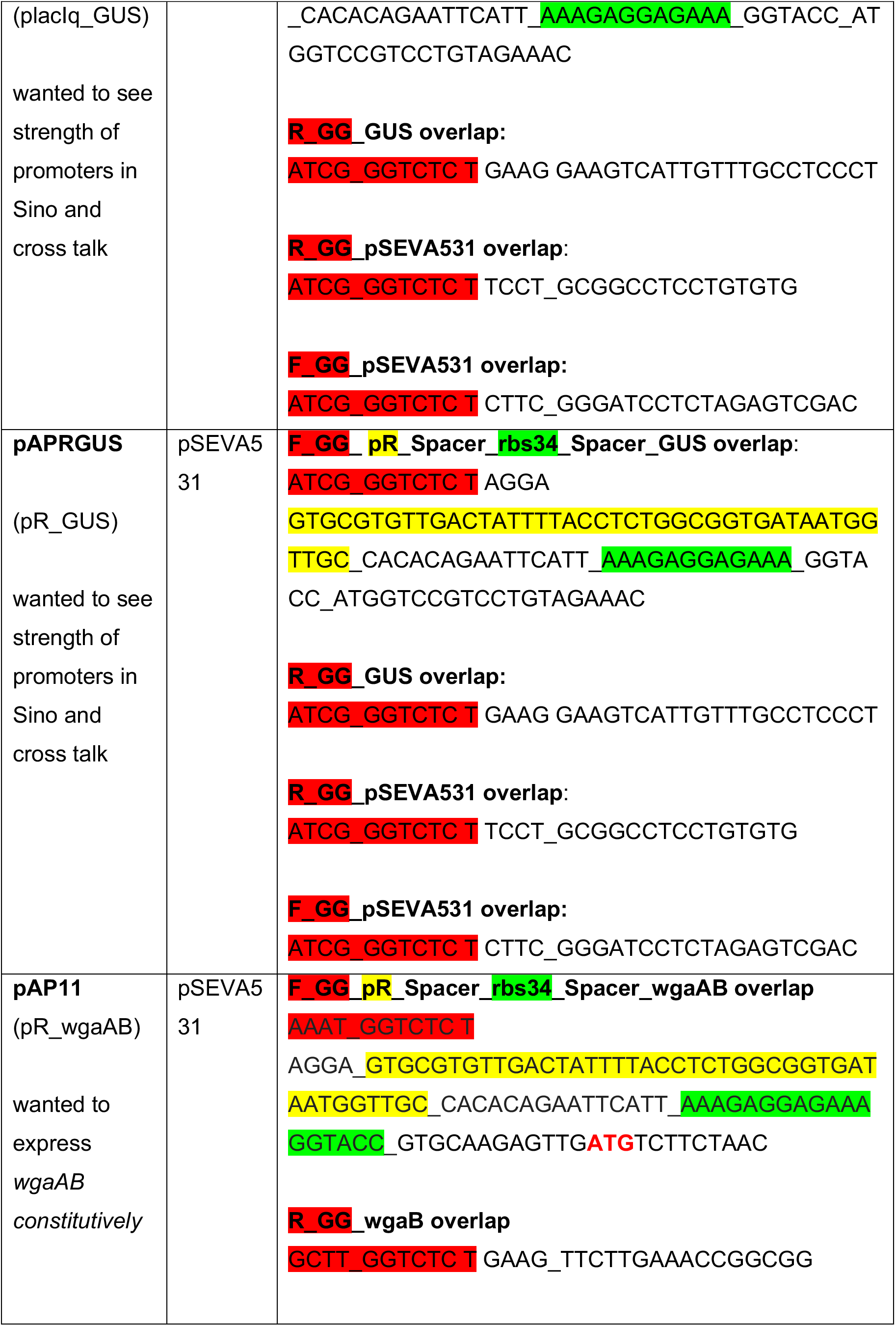
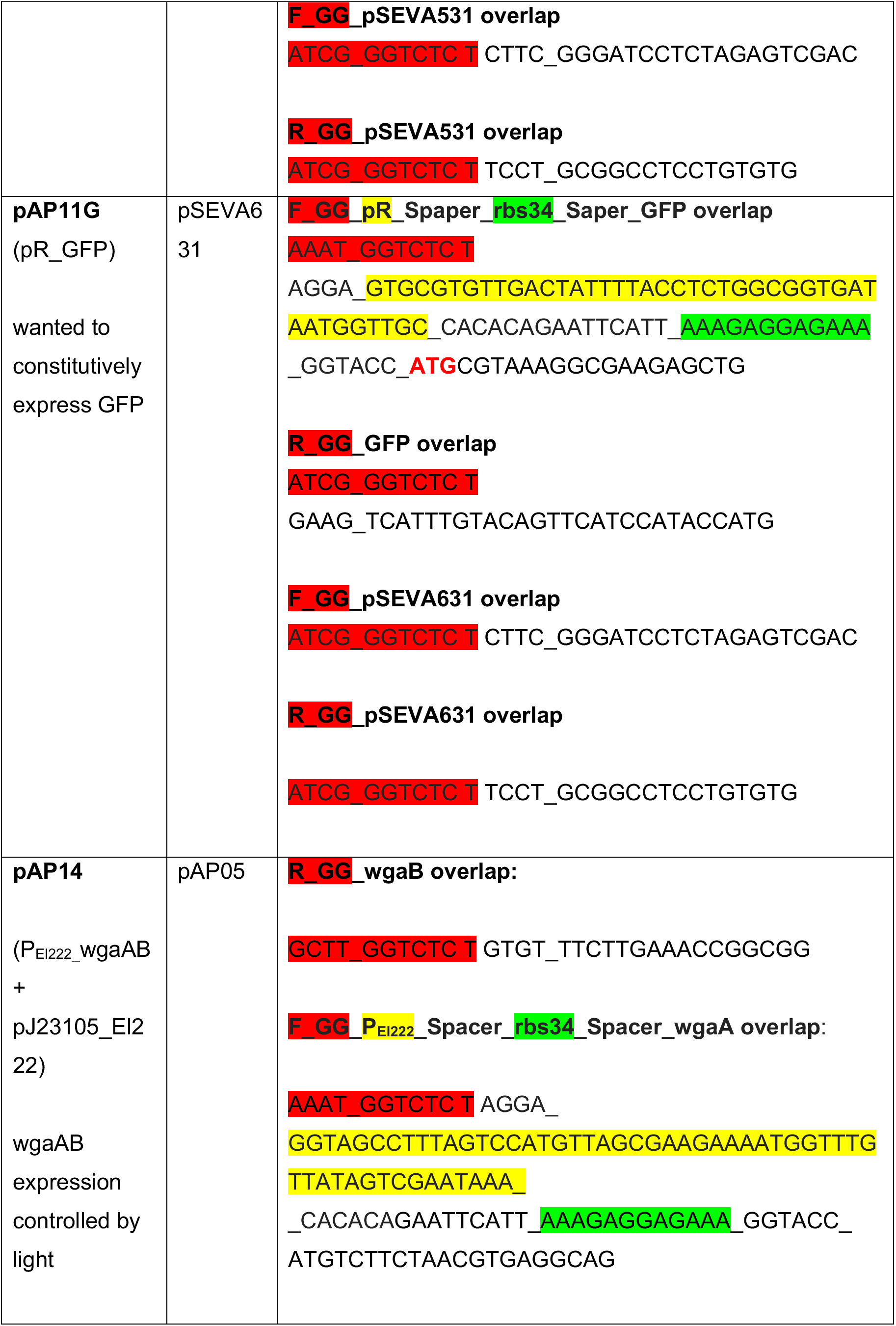
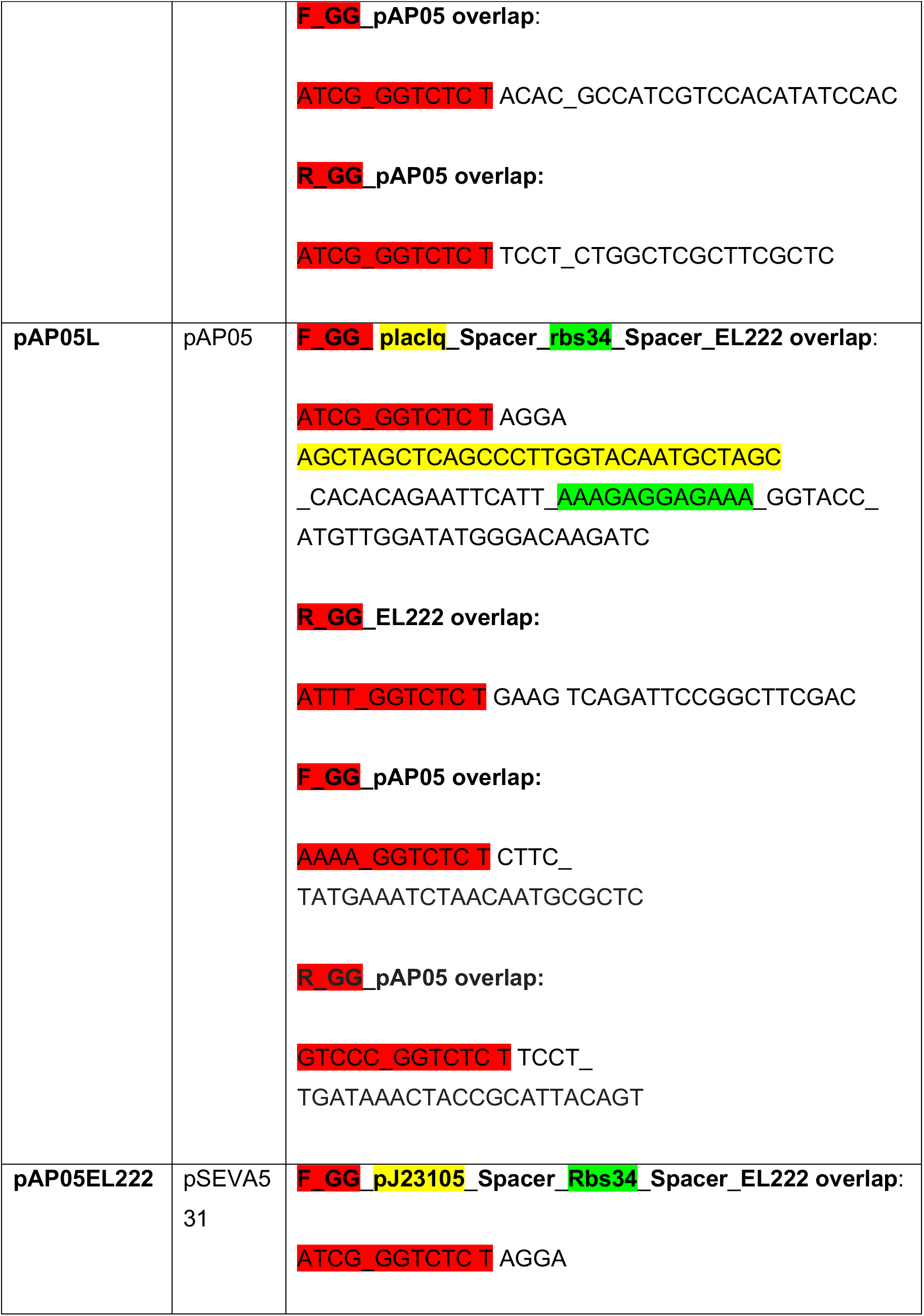
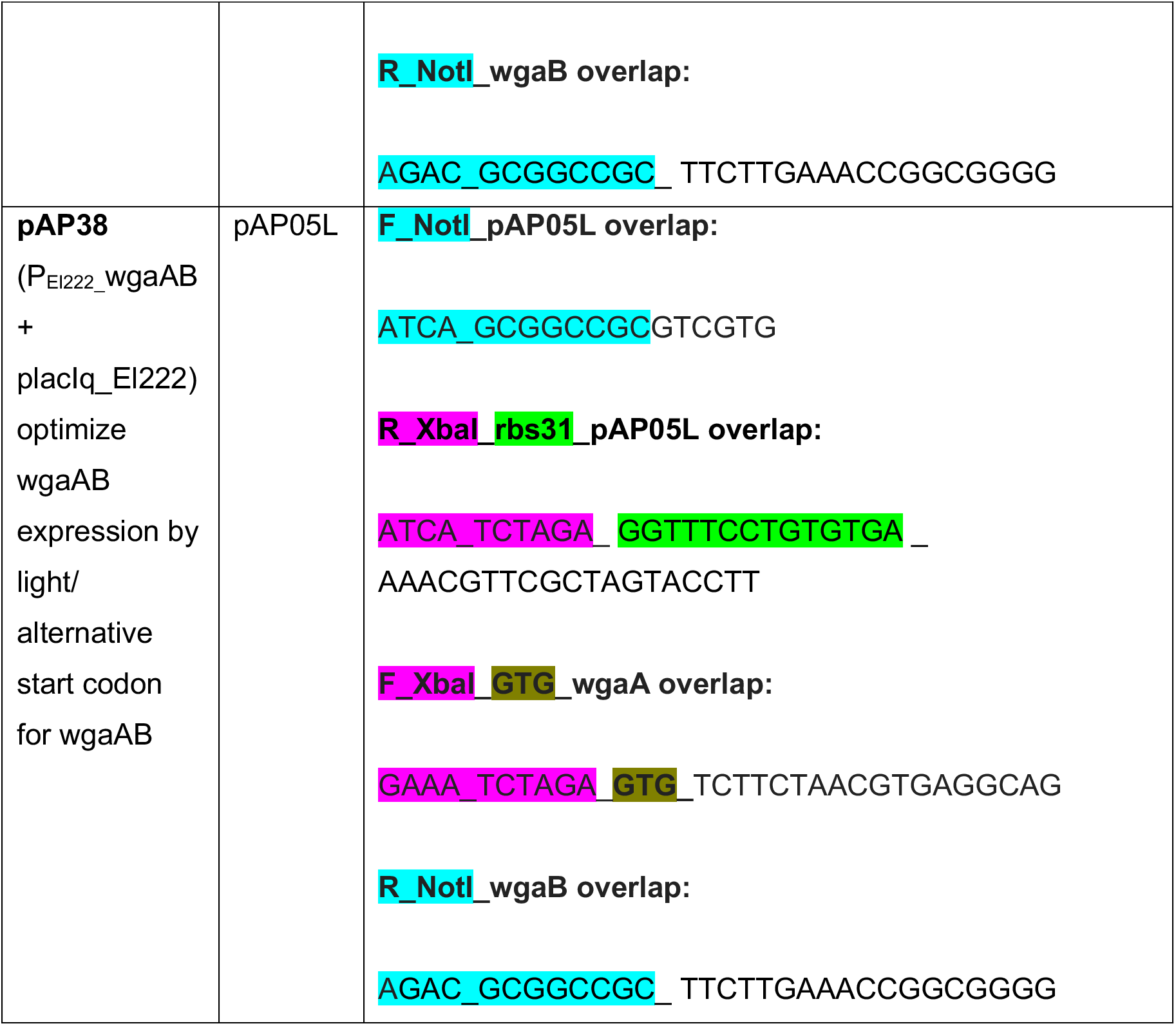
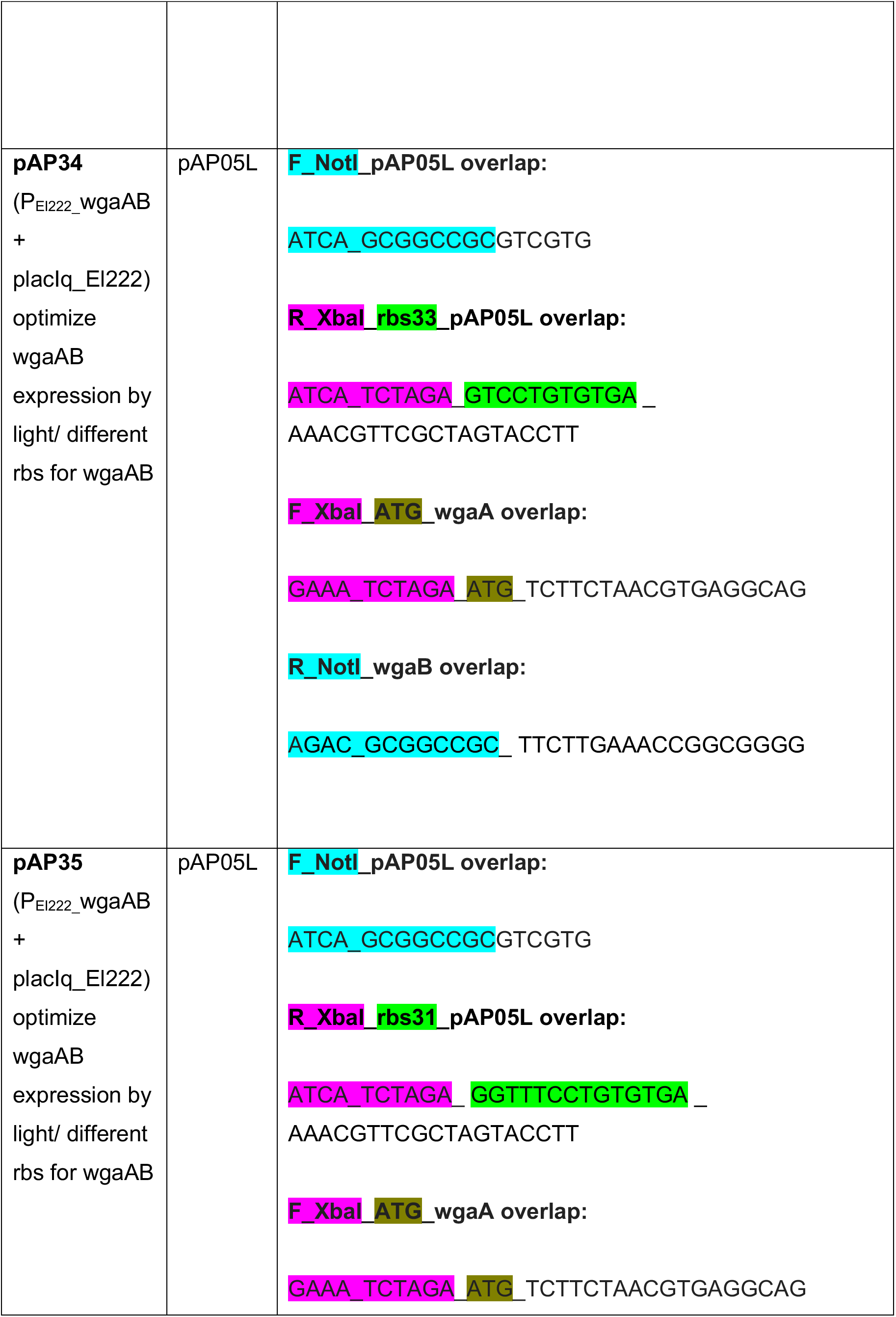
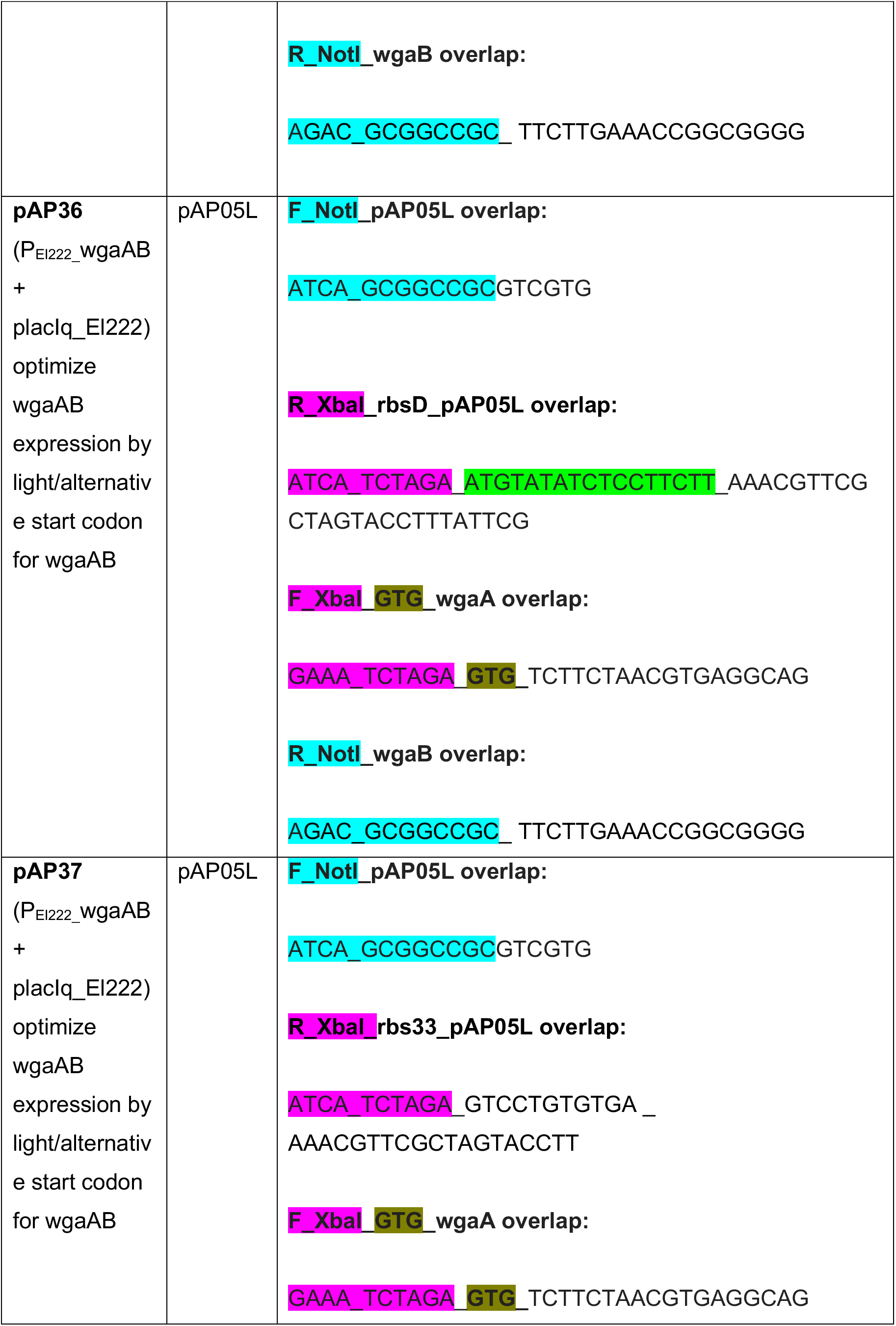
Plasmid map and primers used in this study.

**Supplementary Table 3.** *Sinorhizobium meliloti* strains searched for LOV domain containing proteins.

|  |
| --- |
| *S. meliloti* strain |
| *Sinorhizobium meliloti* 1021 |
| *Sinorhizobium meliloti* 2011 |
| *Sinorhizobium meliloti* AK83 |
| *Sinorhizobium meliloti* B399 |
| *Sinorhizobium meliloti* B401 |
| *Sinorhizobium meliloti* BL225C |
| *Sinorhizobium meliloti* CCMM B554 (FSM-MA) |
| *Sinorhizobium meliloti* GR4 |
| *Sinorhizobium meliloti* HM006 |
| *Sinorhizobium meliloti* KH35c |
| *Sinorhizobium meliloti* KH46 |
| *Sinorhizobium meliloti* M162 |
| *Sinorhizobium meliloti* M270 |
| *Sinorhizobium meliloti* Rm41 |
| *Sinorhizobium meliloti* RU11/001 |
| *Sinorhizobium meliloti* SM11 |
| *Sinorhizobium meliloti* T073 |
| *Sinorhizobium meliloti* USDA1021 |
| *Sinorhizobium meliloti* USDA1106 |
| *Sinorhizobium meliloti* USDA1157 |

Supplementary text related to **Supp. Table 3**.

We searched the proteome of the *Sinorhizobium meliloti* species available at BioCyc Database Collection (biocyc.org), as of 01.05.2020, for potential LOV domain containing proteins (**Supp. Table 3**). Briefly, proteome, open reading frames, of the *S. meliloti* species was scanned for the conserved Asp-Cys-Arg (NCR) tripeptide sequence of LOV domains by using a BLAST search (blast.ncbi.nlm.nih.gov) (13). Subsequently, candidate proteins containing NCR sequence were manually checked for G**X**<u>NCR</u>**Y**MQG (where X = H, Q or K and Y = N, F or L) amino acid sequence (13). Confirming previous reports, we couldn’t find LOV domain containing proteins in *S.meliloti* proteome (14). Not only the LOV domain, but also red-light sensing phytochromes (phy) and blue light sensor BLUF (blue-light sensing using flavin) domains seem to have not evolved in *S. meliloti* (14). However, we cannot exclude the possibility that currently uncharacterized light sensing mechanisms might be present in *S. meliloti* (15).

